# Generative Design of High-Affinity Peptides Using BindCraft

**DOI:** 10.1101/2025.07.23.666285

**Authors:** Mike Filius, Thanasis Patsos, Hugo Minee, Gianluca Turco, Jingming Liu, Monika Gnatzy, Ramon S.M. Rooth, Andy C. H. Liu, Rosa D.T. Ta, Isa H. A. Rijk, Safiya Ziani, Femke J. Boxman, Sebastian J. Pomplun

## Abstract

Discovering high-affinity ligands directly from protein structures remains a key challenge in drug discovery. We applied BindCraft, a structure-guided generative modeling platform, to *de novo* design of peptide ligands for protein interfaces. Originally developed for miniprotein binders, we evaluated its use for shorter peptides (10–20mers) as peptides hold greater synthetic accessibility and therapeutic potential. For the oncoprotein MDM2, BindCraft generated 70 unique peptides; 15 were synthesized, and 7 showed specific binding with nanomolar affinities (K_D_ = 65–165 nM). Competition assays confirmed site-specific binding for the intended target site. For another oncology target, WDR5, peptides were designed for the MLL (WIN) and MYC (WBM) sites. Of the peptides tested for each site, no validated hits were found for the WIN site, but six candidates bound the WBM site with sub-micromolar affinity (K_D_ = 219–650 nM). Based on Bindcraft’s structural prediction of the binding interface, we designed a stapled variant of the best WDR5 binder, improving the potency by 6-fold to a *K*_*D*_ of 39 nM. Overall, our findings establish BindCraft as a powerful and accessible platform for structure-based peptide discovery, with remaining limitations, but with a promising success rate even for challenging targets.

## Introduction

Discovering potent new drugs at the push of a button is a long-standing dream for drug discovery scientists. Structure-based virtual screening over the years has worked on that vision, enabling the *in-silico* exploration of billions of small molecules.^1–4^ Yet despite decades of refinement, virtual hits often show modest binding affinities in the micromolar range, and true hit rates remain frustratingly low – even in recent community benchmarks like the CACHE challenge.^5^

In parallel, computational tools for designing protein-based binders have rapidly advanced. Platforms like Rosetta have generated miniproteins capable of binding target proteins with high potency, resulting in, for example, inhibitors of influenza and SARS-CoV-2.^6–8^ However, these workflows remain technically complex and labor-intensive, often requiring both computational expertise and second stage experimental high throughput screens such as yeast display, to identify validated hits.^6^

A major breakthrough came with AlphaFold, which revolutionized protein structure prediction and was recognized with a Nobel Prize.^9^ Building on this foundation, BindCraft was recently introduced as an AlphaFold-based design tool that generates miniprotein binders directly from a target’s PDB structure. Unlike earlier platforms, BindCraft is remarkably easy to use, requiring minimal computational expertise, and achieves impressive true hit rates (10–100%) across diverse targets.^10^

Despite their elegance as tool compounds, miniproteins face significant hurdles when it comes to translation to therapeutic developments. Their scaffolds raise concerns about immunogenicity, limit bioavailability, and preclude intracellular targeting. Short peptides, by contrast, offer a compelling alternative.^11^ They are synthetically accessible, conformationally flexible, and increasingly drug-like, capable of inhibiting protein–protein interactions while maintaining potential cell permeability and metabolic stability. Recent successes stories such as GLP-1 analogs,^12^ Merck’s PCSK9 peptide inhibitor,^13^ and the macrocyclic KRAS inhibitor LUNA18 underscore their therapeutic potential.^14^

In this study, we evaluate whether BindCraft – originally developed for miniprotein binder design – can be effectively applied to the prediction of short peptide binders (≤20 residues). Even partial success in this domain could offer a cost-effective and scalable alternative to experimental screening methods such as phage or mRNA display. Although several computational approaches for peptide binder prediction have been reported, ^15–17^ few offer the combination of ease-of-use and extensive target validation demonstrated by BindCraft. Nonetheless, applying BindCraft to peptides presents a distinct challenge. Unlike miniproteins, peptides generally lack stable tertiary structures and intramolecular scaffolding, features that often stabilize high affinity binding interfaces.^18^ As such, it is not evident whether the performance observed for miniproteins will translate to this structurally simpler, but therapeutically more promising, class of molecules. Here, we assess the application of Bindcraft to design short peptide binders for the therapeutically relevant oncoprotein MDM2 and for two distinct sites on WDR5. We identify promising initial hit rates among the predicted peptide binders for two out of three of the targets. We further demonstrate that the structural prediction of the binders can be used to enhance their potency through rational chemical modifications, highlighting Bindcraft potential as a practical tool for peptide-based drug discovery. Our study was conducted entirely from the end-user perspective, without software modification or model retraining, ultimately showcasing the usability by typical peptide discovery labs without specialized computational expertise.

## Results

To evaluate the performance of BindCraft in peptide design, we benchmarked the software under conditions representative of typical academic use. BindCraft is freely available, straightforward to install, and offers a user-friendly interface that facilitates rapid deployment even without advanced computational expertise. We downloaded the software from GitHub (https://github.com/martinpacesa/BindCraft) and installed it on a local server equipped with a single GPU. Although BindCraft was originally developed and validated for the generation of small proteins, it includes dedicated filter sets and configurable parameters for peptide-specific applications. We activated these peptide-specific settings and applied additional constraints to generate sequences ranging from 10 to 20 amino acids in length. This configuration provided a practical starting point for assessing the broader applicability of BindCraft to peptide design tasks.

As a benchmarking system, we selected the oncoprotein MDM2, a well-characterized regulator of the tumor suppressor p53 and a key player in cancer biology.^18^ MDM2 binds to the N-terminal transactivation domain of p53, a short α-helical motif, thereby inhibiting its tumor-suppressive function. This interaction has been extensively characterized, and numerous peptide inhibitors have been developed through rational design and high-throughput platforms such as phage display.^18–21^ The abundance of structural data and detailed knowledge of sequence and structural determinants of MDM2 binding make it an ideal model for evaluating peptide generation tools. To initiate peptide design in BindCraft, we uploaded the crystal structure of MDM2 (**Fig. 1a**, PDB: 1YCR). BindCraft allows the user to define single residues or groups of residues on the target protein as design constraints; the software then generates peptide sequences predicted to bind to the selected site. Based on the known p53–MDM2 interface, we selected residues 73-94 as the focal point to guide generation of peptides targeting the p53-binding cleft. Using the peptide-specific configuration described above, we initiated a run requesting 100 candidate sequences. Before exceeding available memory on our server, BindCraft successfully generated 70 unique peptide candidates predicted to bind the MDM2 interface.

**Figure 1:**
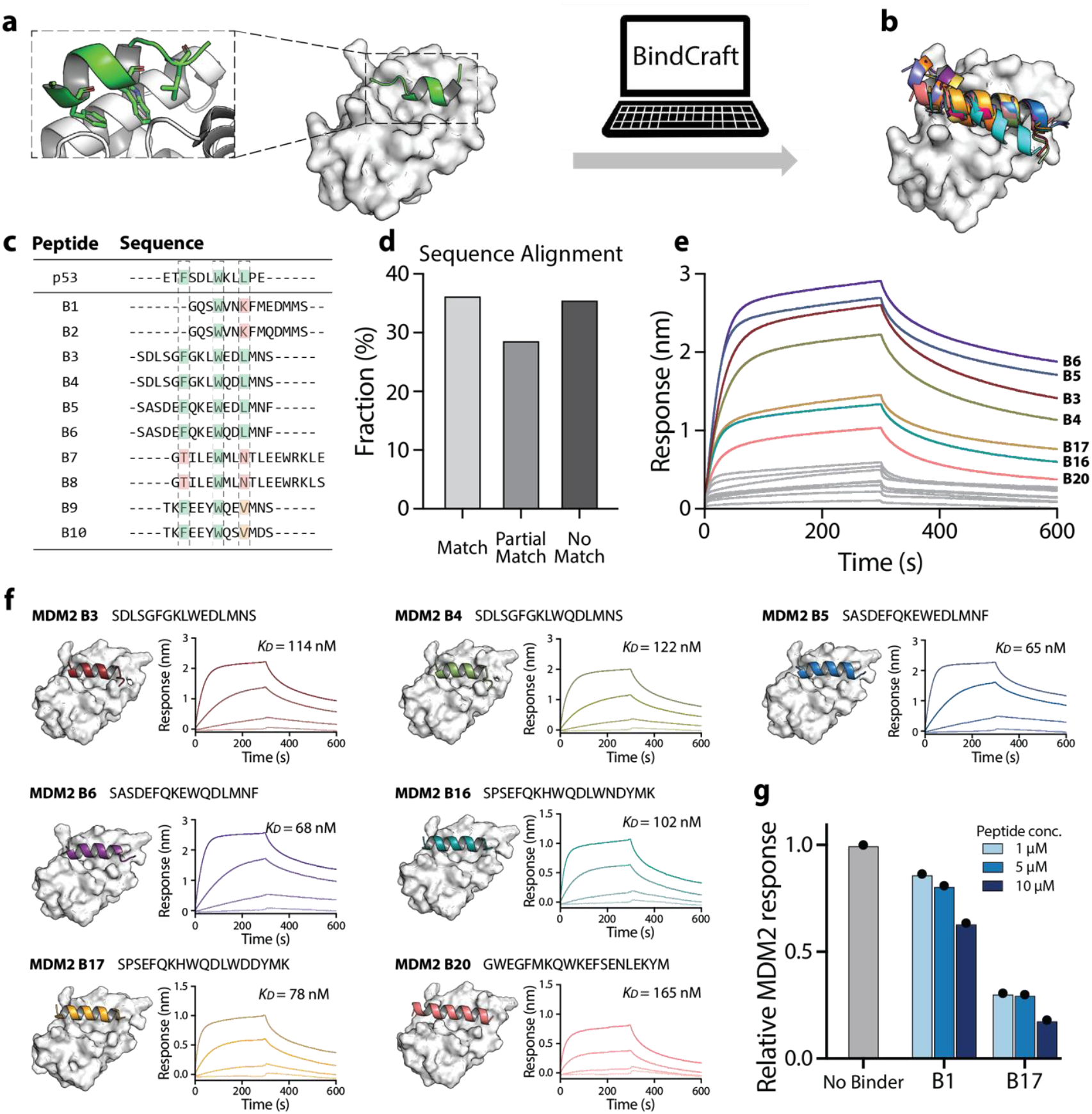
Evaluation of BindCraft’s peptide design capabilities using the MDM2–p53 Interaction. **a)** Crystal structure of MDM2 (white) bound to the N-terminal transactivation domain of p53, a short α-helical motif (green) (PDB: 1YCR). BindCraft was used to design peptides targeting the MDM2–p53 interface by specifying residue 66 of MDM2 as the binding hotspot. **b)** Structures of the top 20 peptide candidates (colored helices) docked at the MDM2 binding site (white). **c)** Sequence alignment of the top 10 designed peptides compared to the reference p53 sequence. We analyzed the presence of the F/W/L hydrophobic hotspot triad: green indicates full triad conservation; orange indicates partial conservation (two of three residues), red indicates mismatch with the triad. **d)** Distribution of hotspot conservation across designed peptides: 36.3% contained the full triad, while 28.7% showed partial conservation, often with alternative hydrophobic residues such as valine (V) or methionine (M). **e)** BLI response curves for the top 15 peptides tested against 1 μM MDM2. High-affinity binders are shown in color; low-affinity or non-binders in grey. **f)** Full binding curves for high-affinity peptides, with dissociation constants (*K*_*D*_) calculated as the average from four different concentrations (1000, 200, 40, 8 nM). **g)** Competition assay to assess specific inhibition of the MDM2–p53 interaction by selected peptides, performed across a range of peptide concentrations (1, 5, 10 μM).

To assess the relevance of the BindCraft-generated peptides, we examined their structural and sequence features in the context of known MDM2-binding motifs. Natural p53-derived peptides, as well as many phage display-optimized variants, adopt an α-helical conformation and rely on a conserved hydrophobic triad (**Fig. 1a**) –phenylalanine (F), tryptophan (W), and leucine (L), at defined spacing – for high-affinity binding to MDM2.^21^ As a first qualitative benchmark, we analyzed the predicted peptides for the presence of these key features. Remarkably, all of the 70 candidate peptides were predicted to adopt α-helical secondary structures (**Fig. 1b**), and 36.3 % contained the full F/W/L hotspot triad (**Fig. 1c and d**). An additional 28.3 % included a partial hotspot motif, featuring two of the three key residues along with other hydrophobic amino acids such as valine (V) or methionine (M), which are also known to contribute to binding in certain engineered variants (**Fig. 1c and d**). Taken together, the structural and sequence characteristics of the generated peptides suggest that BindCraft can produce candidates that closely resemble known MDM2 binders, underscoring its promise for structure-guided peptide design.

To experimentally validate the peptide candidates generated by BindCraft, we selected twenty sequences based on their ranking in the BindCraft output. All peptides were prepared with an N-terminal Lys(biotin) and a β-alanine spacer to facilitate biolayer interferometry (BLI) binding assays. Visual inspection of the peptide–MDM2 interface suggested that this N-terminal modification would not interfere with binding. Fifteen peptides were successfully purified, and screened for binding to MDM2 on the BLI, initially at a single protein concentration (1 μM, **Fig. 1e and Supplementary Table 1**). Seven of the peptides displayed clear association and dissociation kinetics with high-quality fits (R^2^ > 0.95), indicative of specific binding (**Fig 1e**, colored lines). The remaining peptides showed weak or poorly defined binding curves (R^2^ < 0.95), suggesting low affinity or non-specific interactions (**Fig 1e**, grey lines). Full binding curves were subsequently measured for the eight peptides with reliable kinetics, yielding dissociation constants (*K*_*D*_) in the low nanomolar range (65–165 nM, **Fig. 1f**). Notably, six of these seven confirmed binders contained the F/W/L hotspot triad, consistent with its known role in MDM2 recognition.

To confirm the predicted binding site, we performed competition assays against the p53 peptide. To validate their ability to disrupt the native MDM2–p53 interaction, we selected two representative peptides: one strong binder (B17) and one peptide with an unclear binding profile and lacking the full F/W/L hotspot triad (B1). We immobilized a p53-derived peptide on biosensors and measured binding of MDM2 in the presence or absence of varying concentrations of the test peptides. Competitive binding at the p53 site results in a reduced BLI response (**Supplementary Fig. 1)**. Both peptides exhibited dose-dependent inhibition, with B17 showing higher potency (**Fig. 1g**). These results provide strong, albeit indirect evidence that the *de novo* generated peptides engage the predicted binding site on MDM2.

Encouraged by the ability of BindCraft to generate high-affinity binders for MDM2, we next evaluated its performance on a second biologically and therapeutically relevant target: WDR5.^22^ WDR5 is a core component of multiple chromatin-modifying complexes and plays a central role in epigenetic regulation, stem cell maintenance, and oncogenesis.^22,23^ It is also implicated in transcriptional activation of oncogenes such as *MYC*, making it an emerging target in cancer therapy.^24,25^ Structurally, WDR5 presents two well-characterized and pharmacologically interesting binding sites: the WIN site, which binds peptides derived from the MLL (mixed lineage leukemia) protein,^26^ and a second site that engages the MYC transactivation domain.^27^ To test whether BindCraft could generate distinct peptide ligands for each site, we designed two separate prediction runs. For the WIN site, we uploaded the WDR5 structure (PDB: 6DY7) and selected residue F133 as the design anchor; BindCraft successfully generated 100 peptide candidates targeting this region. In a second run, we selected residue L240 to focus on the MYC-binding interface, resulting in a separate set of 100 predicted peptides. Interestingly, all peptides generated for both sites were predicted to adopt α-helical conformations (**Fig. 2a and b**), even though the natural MLL and MYC peptides bind WDR5 as largely unstructured loops rather than helices (**Supplementary Fig. 2**).

**Figure 2:**
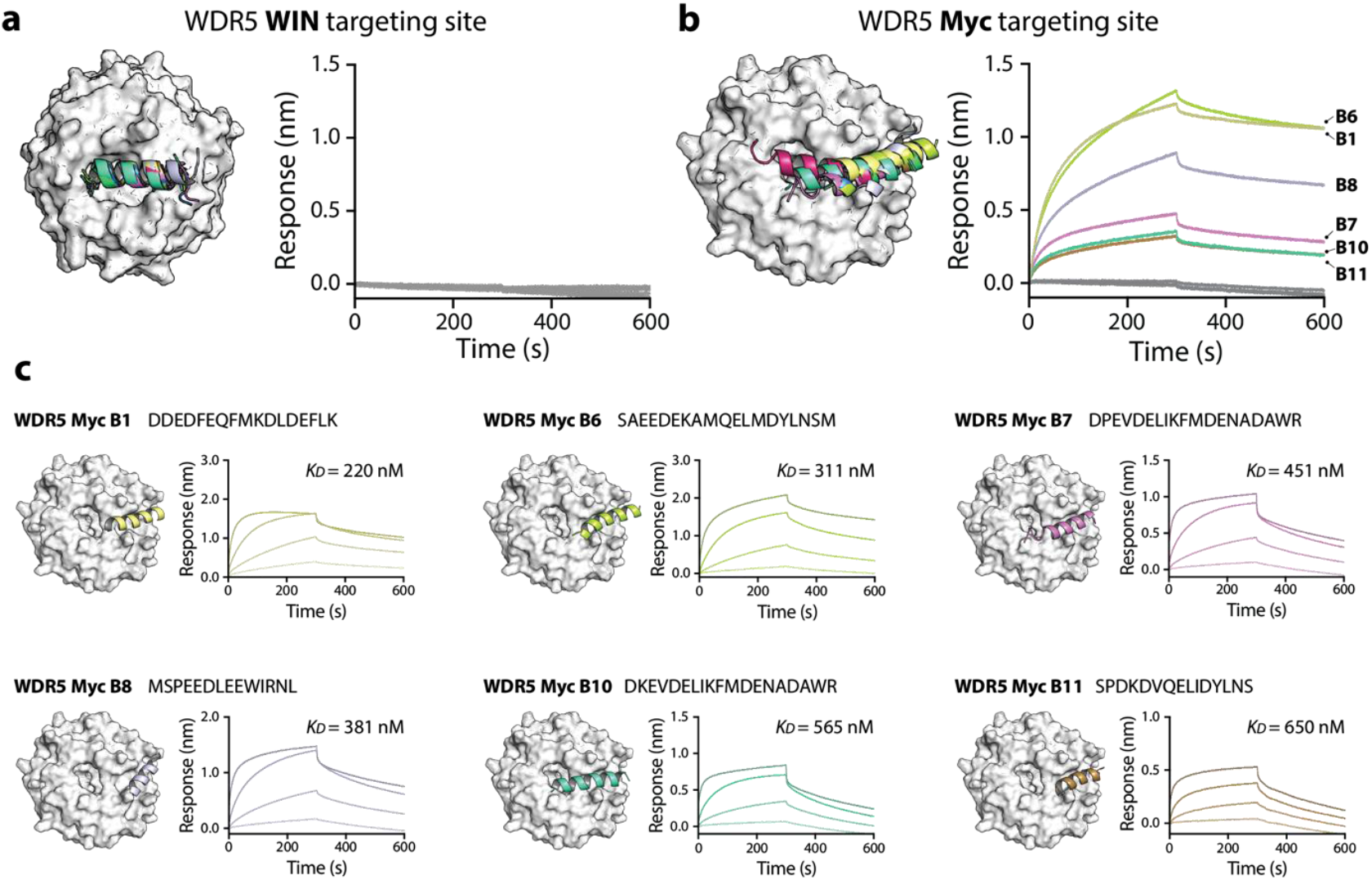
de novo design and characterization of peptides binders to WDR5. **a)** Crystal structure of WDR5 (white; PDB:6DY7) with predicted peptides targeting the MLL (WIN) binding site shown as colored helices. **Right:** Binding affinities of the top 10 predicted peptides assessed via BLI; none showed detectable binding to WDR5. **b)** Crystal structure of WDR5 (white; PDB: 6DY7) with predicted peptides targeting the Myc binding site (colored helices). **Right:** BLI-based affinity measurements for the Myc-site binders; 6 out of 9 peptides exhibited moderate binding to WDR5. **c)** Full binding curves for high-affinity Myc-site peptides. Dissociation constants (*K*_*D*_) were determined as the average from four different concentrations (10, 2, 0.4, 0.08 μM).

Given the high success rate observed for MDM2, we prioritized the top ten peptides from each WDR5 peptide set for experimental validation. Peptides were synthesized with the same N-terminal Lys(biotin) and β-alanine linker to enable biolayer interferometry (BLI) assays. Binding was initially screened at a single concentration of 1 μM WDR5, following the same protocol as before. Interestingly, none of the successfully purified peptides targeting the MLL (WIN) site showed detectable binding (**Fig. 2a, Supplementary Table**). In contrast, six out of nine peptides designed for the MYC-binding site exhibited clear association and dissociation kinetics (**Fig. 2b**). Full kinetic analysis of these six peptides yielded dissociation constants (*K*_*D*_) ranging from 219 to 650 nM, confirming moderate affinity interactions at this challenging target interface. While these peptides are not predicted to adopt the same conformation as the natural MYC peptide when bound to WDR5, they share the characteristic of containing multiple negatively charged residues, which may contribute to their binding affinity.

Bindcraft’s structural predictions enable the identification of optimal peptide stapling positions, obviating the need for experimental mutation scanning. Peptide staples reinforce α-helical conformations by linking side chains on the same face of the helix, typically one or two turns apart (i, i+4 or i, i+7).^28,29^ This conformational locking can enhance peptide binding affinity by stabilizing the bioactive structure, and also improve proteolytic stability and sometimes cellular permeability.^19,30^ Identifying stapling sites that do not disrupt the peptide– protein interface is critical. Traditionally, this requires structural data. In the absence of such data, alanine scanning is often used to identify mutable residues and potential stapling pairs— an approach that is experimentally intensive. By contrast, Bindcraft’s outputs can be directly applied to the design of stapled peptide variants, bypassing the need for further experimental input. As a case study, we analyzed the WDR Myc_B1 peptide, predicted to bind WDR5 in an α-helical conformation – making it a strong candidate for stapling. The structural prediction clearly shows the largely hydrophobic binding interface, and backside residues that do not appear to be involved in protein binding (**Fig. 3a and 3b**). We selected two solvent-exposed residues on the helix’s backside, Glu6 and Lys10, and mutated them to cysteines to enable meta-xylene stapling.^31^ The resulting stapled peptide exhibited a dissociation constant (*K*_*D*_) of 39 nM, a ∼6-fold improvement over the unstapled parent peptide. Finally, we assessed the ability for the stapled Myc_B1 peptide to disrupt the native Myc-WDR5 interaction. We immobilized the WDR5 binding MYC fragment and measured binding of WDR5 in the presence or absence of varying concentrations of the test peptides. We observed Concentration dependent inhibition is observed for the stapled Myc_B1 variant reaching a maximum inhibition of 57 % at 20 μM, while the native Myc_B1 has a maximum inhibition of only 25 % at 20 μM (**Fig. 3f and Supplementary Fig. 3**). Altogether, these results highlight Bindcraft’s advantage over structure-agnostic platforms like phage display: its structural predictions of validated binders can be immediately leveraged for rational peptide optimization.

**Figure 3:**
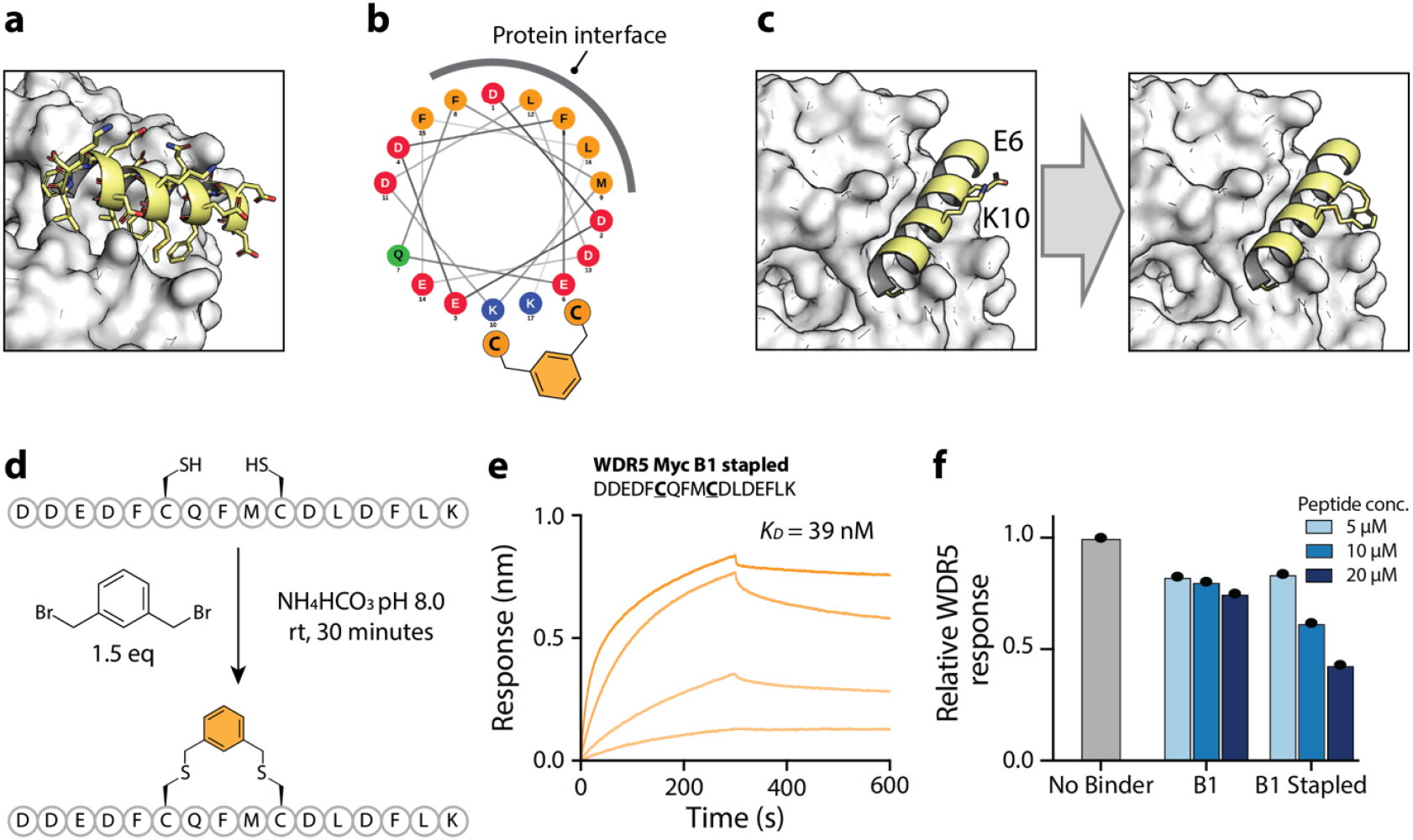
de novo design and characterization of peptides binders to WDR5. **a)** Structural prediction of Myc_B1 with WDR5 showing the residues that are involved in protein binding. **b)** wheel projection of Myc_B1 revealing the largely hydrophobic binding residues involved in protein binding. Figure made with NetWheels <u>(NetWheels: Peptides Helical Wheel and Net projections maker</u>). **c)** two solvent-exposed residues on the helix’s backside, Glu6 and Lys10, and mutated them to cysteines to enable meta-xylene stapling. **d)** cysteines were introduced in an *i, i+4* configuration and stapled with 1,3-bis(bromomethyl)-benzene. **e)** Full binding curves for the stapled Myc_B1 binder. Dissociation constants (*K*_*D*_) were determined as the average from four different concentrations (2.5, 0.5, 0.1, 0.2 μM). **f)** Competition assay to assess specific inhibition of the WDR5–Myc interaction by the native Myc_B1 peptide vs the stapled Myc_B1 variant, performed across a range of peptide concentrations. Concentration dependent inhibition is observed for the stapled Myc_B1 variant reaching a maximum inhibition of 57% at 20 μM, while the native Myc_B1 has a maximum inhibition of 25% at 20 μM.6

## Discussion and Conclusion

This study demonstrates the feasibility of using the generative modeling platform BindCraft – originally developed for miniprotein design – as a powerful tool for structure-based discovery of short peptide binders. Peptides, in contrast to miniproteins, offer distinct advantages in terms of chemical accessibility, and drug-likeness, and are thus a highly relevant molecular format for next-generation therapeutics. While AlphaFold-based approaches have proven highly effective for peptide–protein docking and structural prediction,^32,33^ the *de novo* generation of functional peptide ligands is still an emerging area.

We assessed the BindCraft platform from the perspective of typical end users, without modifying code or model parameters. Using the publicly available version of BindCraft “off the shelf,” we applied it to the discovery of peptide ligands for the oncoprotein MDM2 and for two distinct binding sites on the chromatin-associated protein WDR5. For two out of these three targets - MDM2 and the MYC-binding site of WDR5 – we identified validated binders with nanomolar to sub-micromolar affinities, demonstrating that BindCraft can generate functional peptide ligands from structural input alone. We later extended this approach to the PD-1/PD-L1 immune checkpoint interface; however, none of the predicted peptides resulted in detectable binding in BLI assays (data not shown). Further investigations into target-specific success factors and optimization strategies are ongoing.

In a striking example, we demonstrated the advantage of using chemically accessible peptides for rapid post-discovery optimization guided by predicted structures. Importantly, we show that BindCraft not only generates functional peptide sequences but also provides reliable structural models that can inform chemical modification. This was exemplified by a stapling experiment in which structure-guided residue selection led to a six-fold improvement in binding affinity, an outcome that underscores the practical utility of generative models for downstream optimization. Such rational enhancements are typically out of reach for screening-based methods like phage or mRNA display, which lack intrinsic structural information for validated binders.

Interestingly, all peptides generated by BindCraft in this study were predicted to adopt α-helical conformations. While this is consistent with the MDM2–p53 interface – where the native binding partner is a well-defined helix – it is less intuitive for the two WDR5 binding sites. Both the MLL and MYC peptides engage WDR5 as largely unstructured loops or random coils in their native complexes. The bias toward helical structures likely reflects an inherent tendency of generative models to favor stable, well-defined secondary structures, such as α-helices, which are easier to model and are more likely to receive confident predictions from structure-prediction engines like AlphaFold. This preference may improve design success rates overall but could limit access to non-helical binding modes, particularly for targets with flexible or irregular interfaces.

While BindCraft exhibited limitations for certain protein–protein interfaces – such as PD-1/PD-L1 and the WDR5 WIN site, where no validated binders were obtained – it successfully produced high-affinity, site-specific peptide ligands for MDM2 and the WDR5 MYC site. In both successful cases, the hit rates were remarkably high, especially considering that only a modest number of candidate peptides were synthesized, highlighting the feasibility of this approach for research groups with limited screening capacity. A key advantage over display-based methods is that the user defines the binding site, enabling the design of targeted inhibitors for specific interfaces rather than relying on selection from a diverse but uncontrolled pool. Taken together, our results suggest that tools like BindCraft are reaching a level of maturity where they can serve as realistic and accessible alternatives to traditional display technologies for initial hit discovery in peptide-based drug development.

## Acknowledgements

M.F. and S.J.P. acknowledge funding from NWO (OCENW.M.23.152). The Pomplun Lab gratefully acknowledges financial support by Mr. H. J. M. Roels through a donation to the Oncode Institute and KWF’s financial support of the Oncode Institute.

## Author Contributions

Conceptualization: M.F. and S.J.P.

Synthesis: M.F., T.P., H.M., G.T., J.L., M.G., R.S.J.R., A.C.H.L, R.D.T.T., I.H.A.R., S.Z., and F.J.B.

Kinetic data acquisition and analysis: M.F., and S.J.P. Writing – original draft: M.F. and S.J.P.

Writing – review & editing: M.F., T.P., H.M., G.T., J.L., M.G., R.S.J.R., A.C.H.L, R.D.T.T., I.H.A.R., S.Z., F.J.B., and S.J.P.

## Data and Materials Availability

All the data required to support the conclusions in the paper are included in the paper itself and/or the Supplementary Materials. Any additional data are available from the corresponding author upon request.

## Materials and Methods

### BindCraft design settings for protein targets

The BindCraft software was installed on an in-house server following the instructions at https://github.com/martinpacesa/BindCraft. Peptide designs for the targets described in the Results section were generated using the ‘peptide_filters_relaxed’ filter set in combination with the ‘peptide_3stage_multimer’ advanced settings provided by the BindCraft software. Cysteine residues were excluded from all peptide designs. Details of the input structures and hotspot designations are provided in **Supplementary Table 4**. For experimental validation, the top 10 peptide designs were selected for each target, except for MDM2, for which the top 20 designs were tested.

### Peptide synthesis

Peptides were synthesized on preloaded ProTide Rink amide resin (CEM, R003; 83 mg, 0.6 mmol/g). The resin was swollen in DMF, deprotected with 20% piperidine in DMF (5 min), and washed with DMF (5×). Fmoc-protected amino acids (Fmoc-AA-OH, Sigma-Aldrich) were preactivated at 0.275 M (5.5 eq) in 0.25 M HATU (5.0 eq, Fluorochem) in DMF. DIPEA (100 μL, Fluorochem) was added to 1 mL of the activated amino acid solution prior to coupling. Each coupling was performed for 15 min at room temperature, followed by DMF washes (5×) and Fmoc deprotection (5 min, 20% piperidine in DMF). This cycle was repeated until full sequence assembly, including a β-alanine spacer. For C-terminal biotinylation, Fmoc-Lys(biotin)-OH (BLDPharm) was coupled at 0.2 M in DMF with HATU and DIPEA, with an extended reaction time of 2 h. After final Fmoc removal, N-terminal acetylation was performed using an acetylation cocktail (acetic anhydride:DIPEA:DMF, 1:1:8) for 30 min. The resin was washed with DMF (5×) and Et2O (3×), then air-dried. Peptides were cleaved and globally deprotected using 2 mL of cleavage cocktail (87.5% TFA, 5% Milli-Q water, 5% 1,2-ethanedithiol, 2% triisopropylsilane) for 3 h at room temperature. Crude peptides were precipitated in ice-cold diethyl ether and redissolved in water/acetonitrile for purification by reverse-phase HPLC. Peptide identity was confirmed by mass spectrometry. Corresponding LC and MS traces are shown in **Supplementary Figures 4-37**.

### Stapling (only when successful)

For the WDR5-Myc B1 stapling reaction, we synthesized a peptide containing two cysteines in an *i, i+4* configuration (**Supplementary Table 5**). The cysteine-containing peptide was synthesized, cleaved, biotinylated, and purified as described in the *Peptide Synthesis* section. Stapling was performed at a peptide concentration of 1 mM using 1.5 equivalents of 1,3-bis(bromomethyl)-benzene (Sigma-Aldrich) in stapling buffer (50mM NH_4_HCO_3_ (pH 8.0), 25% Acetonitrile, in Mill-Q water) for 30 minutes at room temperature. Reaction completion was confirmed by mass spectrometry, after which the stapled peptide was immediately purified by reverse-phase HPLC. The corresponding LC and MS trace can be found in **Supplementary Figures 38 and 39**.

### Biolayer interferometry (BLI)

The BLI experiments were performed in 96 well plates (GreinerBio-One, polypropylene, flat-bottom, chimney well) using an Octet R4 system (SATORIUS). Streptavidin biosensors were functionalized by incubating them for 60 s in 200 μL of 1 μM purified, biotinylated peptide prepared in kinetic buffer (1x PBS, 0.02% Tween-20, 1mg/mL BSA). Subsequently, a baseline was established by immersing the biosensors in kinetic buffer for 60 s. The sensors were then incubated in 200 μL of protein sample for 300 s to measure the association phase, followed by a 300 s incubation in kinetic buffer to assess the dissociation phase. The measurements were performed at room temperature. The kinetic rates and dissociation constant (*K*_*D*_) were determined by fitting a 1:1 kinetic model using the Octet R4 analysis software.

### WDR5 Expression and Purification

The truncated WDR5 (amino acids 22–334) was cloned into a pET vector with a 6xHis-SUMO tag fused at the N-terminus. The WDR5 plasmid was then transformed into E. coli BL21 (DE3) cells. The cells were cultured in Luria–Bertani medium at 37 °C. When the optical density at 600 nm (OD600) reached 0.8, the temperature was lowered to 25 °C. Protein expression was induced by adding 1 mM isopropyl-β-D-thiogalactoside (IPTG), and the incubation continued for 16 hours at this temperature. The cells were harvested by centrifugation, resuspended in lysis buffer (20 mM Tris-HCl, 500 mM NaCl, pH 8.0), and then lysed using a homogenizer (FPG12800). The lysate was cleared by centrifugation, and the supernatant was collected. The protein was then bound to a nickel affinity column (HisTrap™, Cytiva) using an ÄKTA system and eluted using an imidazole gradient. The purified protein was verified by SDS-PAGE and concentrated using a 10K molecular weight cut-off centrifugal concentrator. The concentration was determined using a NanoDrop spectrophotometer.

