## Supplementary Information for "Generative Design of High-Affinity Peptides Using BindCraft"

### Generative Design of High-Affinity Peptide Binders Using BindCraft


| | |
|---|---|
| Supplementary Table 1 | BLI screening for MDM2 binders at single concentration. |
| Supplementary Fig. 1 | Competition assay for MDM2 with binders B1 and B17 |
| Supplementary Fig. 2 | Crystal structures for WDR5 in complex with the MLL and Myc binding peptide |
| Supplementary Table 2 | BLI screening for WDR5 WIN binders at single concentration |
| Supplementary Table 3 | BLI screening for WDR5 Myc binders at single concentration |
| Supplementary Fig. 3 | Competition assay for WDR5 with binders B1 and B1 stapled. |
| Supplementary Table 4 | Overview of BindCraft input settings used for different protein targets. |
| Supplementary Table 5 | WDR5 Myc B1 stapled sequence. |
| Supplementary Figs. 4-19 | LC-MS Analysis for MDM2 peptides |
| Supplementary Figs. 20-32 | LC-MS Analysis for WDR5 peptides |
| Supplementary Fig. 33 | LC-MS Analysis for Myc-peptide-motif for WDR5 competition assay |
| Supplementary Fig. 34 | LC-MS Analysis for p53-peptide-motif for MDM2 competition assay |
| Supplementary Figs. 35 and 36 | LC-MS Analysis for MDM2 B1 and B17 used for competition assay |
| Supplementary Figs. 37 | LC-MS Analysis for WDR5_Myc_B1 used for competition assay |
| Supplementary Figs. 38 and 39 | LC-MS Analysis for WDR5_Myc_B1 used for stapling and competition assay |

**Supplementary Table 1:** BLI screening for MDM2 binders at 1 µM of MDM2.

| Binder | KD (M) | KD Error | ka (1/Ms) | ka Error | kdis (1/s) | kdis Error | Full R^2 |
|---|---|---|---|---|---|---|---|
| MDM2_B1 | 3.428E-07 | 1.051E-07 | 1.141E04 | 2.459E03 | 3.913E-03 | 8.534E-04 | 0.8456 |
| MDM2_B11 | 2.500E-07 | 4.357E-08 | 3.542E04 | 4.051E03 | 8.856E-03 | 1.164E-03 | 0.9237 |
| MDM2_B12 | 2.756E-07 | 6.971E-08 | 1.672E04 | 2.920E03 | 4.609E-03 | 8.432E-04 | 0.8588 |
| MDM2_B13 | 2.044E-07 | 3.998E-08 | 2.994E04 | 4.196E03 | 6.119E-03 | 8.348E-04 | 0.8879 |
| MDM2_B14 | 4.956E-07 | 1.713E-07 | 8.154E03 | 2.008E03 | 4.041E-03 | 9.801E-04 | 0.8403 |
| MDM2_B16 | 4.331E-07 | 3.445E-08 | 1.801E04 | 7.948E02 | 7.800E-03 | 5.162E-04 | 0.9853 |
| MDM2_B17 | 3.044E-07 | 2.701E-08 | 1.873E04 | 1.115E03 | 5.702E-03 | 3.753E-04 | 0.9729 |
| MDM2_B18 | 3.340E-07 | 8.472E-08 | 1.439E04 | 2.557E03 | 4.806E-03 | 8.696E-04 | 0.8752 |
| MDM2_B2 | 6.174E-07 | 1.581E-07 | 9.642E03 | 1.757E03 | 5.953E-03 | 1.071E-03 | 0.9006 |
| MDM2_B20 | 7.359E-07 | 1.052E-07 | 1.316E04 | 9.160E02 | 9.687E-03 | 1.210E-03 | 0.9763 |
| MDM2_B3 | 1.175E-06 | 1.781E-07 | 8.517E03 | 1.926E02 | 1.000E-02 | 1.500E-03 | 0.9932 |
| MDM2_B4 | 9.647E-07 | 1.048E-07 | 9.395E03 | 2.304E02 | 9.064E-03 | 9.589E-04 | 0.9913 |
| MDM2_B5 | 8.376E-07 | 1.249E-07 | 1.165E04 | 2.062E02 | 9.757E-03 | 1.444E-03 | 0.9901 |
| MDM2_B6 | 2.639E-06 | 5.577E-08 | 9.427E03 | 5.397E01 | 2.488E-02 | 5.061E-04 | 0.9962 |
| MDM2_B8 | 7.830E-07 | 1.598E-07 | 8.044E03 | 9.689E02 | 6.299E-03 | 1.038E-03 | 0.9296 |

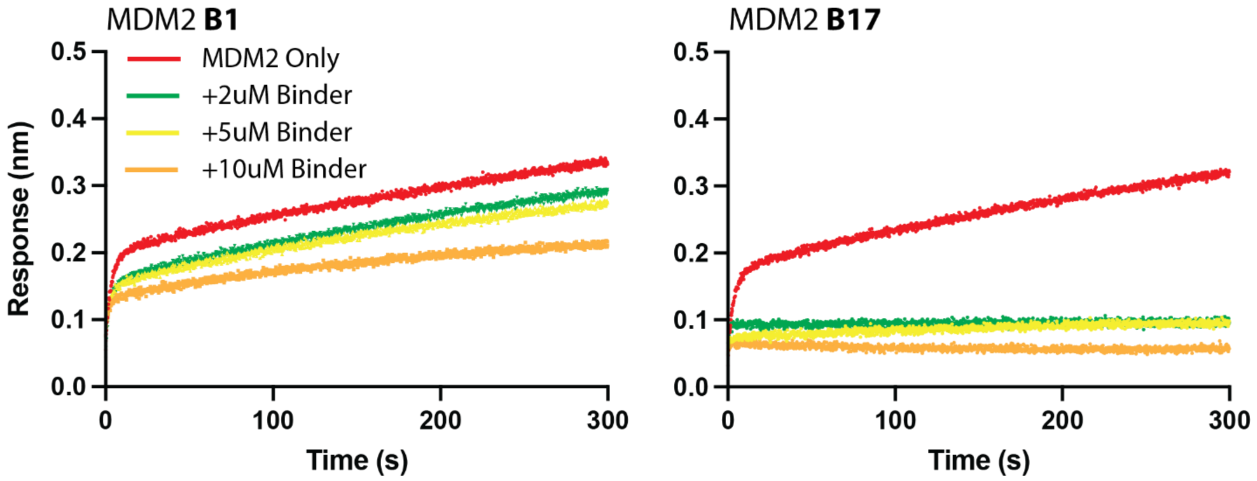


**Supplementary Figure 1: Competition assay for MDM2 with binders B1 and B17.** Competition analysis (BLI association) of two predicted binders (B1 and B17) to MDM2-p53. P53 immobilized on BLI tips was dipped into solutions containing 1 µM MDM2 and different concentrations of the binders (2, 5, and 10 µM). Increasing the concentration of the predicted binders (B1 and B17) in solution causes occupancy of the MDM2-p53 binding pocket and results in a concentration-dependent decrease in BLI response.

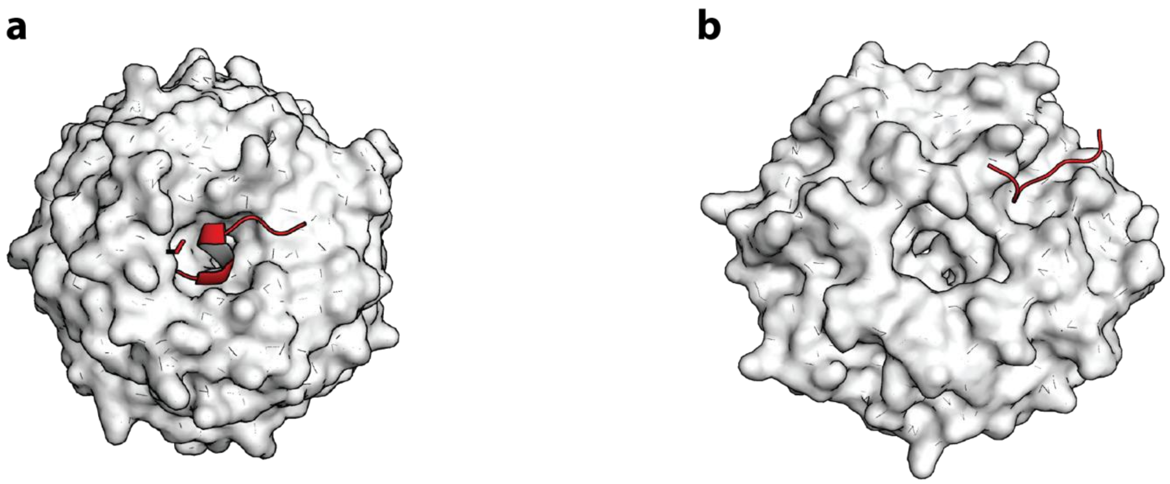


**Supplementary Figure 2:** Crystal structures for WDR5 in complex with the MLL binding motif (panel a, PDB: 3UVM) and Myc binding motif (panel b, PDB: 4Y7R). peptides are highlighted in red, WDR5 crystal structure in white.

**Supplementary Table 2:** BLI screening for WDR5 WIN binders at 2 µM of WDR5.

| Sample ID | Response | KD (M) | KD Error | ka (1/Ms) | ka Error | kdis (1/s) | kdis Error |
|---|---|---|---|---|---|---|---|
| Win_B1 | -1.780E-02 | 1.000E-07 | 0.000E00 | 1.000E04 | 0.000E00 | 1.000E-03 | 0.000E00 |
| Win_B8 | -3.945E-02 | 1.000E-07 | 0.000E00 | 1.000E04 | 0.000E00 | 1.000E-03 | 0.000E00 |
| Win_B2 | -1.082E-02 | 1.000E-07 | 0.000E00 | 1.000E04 | 0.000E00 | 1.000E-03 | 0.000E00 |
| Win_B9 | -4.241E-02 | 1.000E-07 | 0.000E00 | 1.000E04 | 0.000E00 | 1.000E-03 | 0.000E00 |
| Win_B5 | -3.939E-02 | 1.000E-07 | 0.000E00 | 1.000E04 | 0.000E00 | 1.000E-03 | 0.000E00 |
| Win_B10 | -3.857E-02 | 1.000E-07 | 0.000E00 | 1.000E04 | 0.000E00 | 1.000E-03 | 0.000E00 |
| Win_B6 | -4.174E-02 | 1.000E-07 | 0.000E00 | 1.000E04 | 0.000E00 | 1.000E-03 | 0.000E00 |
| Win_B11 | -2.580E-02 | 1.000E-07 | 0.000E00 | 1.000E04 | 0.000E00 | 1.000E-03 | 0.000E00 |

**Supplementary Table 3:** BLI screening for WDR5 Myc binders at 2 μM of WDR5.

| Sample ID | Response | KD (M) | KD Error | ka (1/Ms) | ka Error | kdis (1/s) | kdis Error | Full R^2 |
|---|---|---|---|---|---|---|---|---|
| Myc_B1 | 1.2214 | 4.396E-08 | 6.287E-10 | 8.282E03 | 3.012E01 | 3.641E-04 | 5.036E-06 | 0.9858 |
| Myc_B8 | 0.8834 | 1.045E-07 | 1.567E-09 | 6.985E03 | 4.794E01 | 7.303E-04 | 9.731E-06 | 0.9532 |
| Myc_B3 | -9.846E-03 | 1.000E-07 | 0.000E00 | 1.000E04 | 0.000E00 | 1.000E-03 | 0.000E00 | |
| Myc_B10 | 0.3503 | 2.040E-07 | 2.815E-09 | 9.039E03 | 9.548E01 | 1.844E-03 | 1.638E-05 | 0.902 |
| Myc_B5 | 0.0129 | 2.598E-04 | 4.592E-02 | 1.930E03 | 3.411E05 | 5.014E-01 | 4.768E-01 | 0.1212 |
| Myc_B11 | 0.3188 | 1.869E-07 | 2.367E-09 | 7.889E03 | 6.984E01 | 1.475E-03 | 1.335E-05 | 0.9241 |
| Myc_B6 | 1.3049 | 9.236E-08 | 1.278E-09 | 6.037E03 | 3.147E01 | 5.576E-04 | 7.150E-06 | 0.9779 |
| Myc_B12 | -1.519E-02 | 1.000E-07 | 0.000E00 | 1.000E04 | 0.000E00 | 1.000E-03 | 0.000E00 | |
| Myc_B7 | 0.4711 | 1.455E-07 | 1.938E-09 | 1.068E04 | 1.030E02 | 1.553E-03 | 1.427E-05 | 0.9054 |

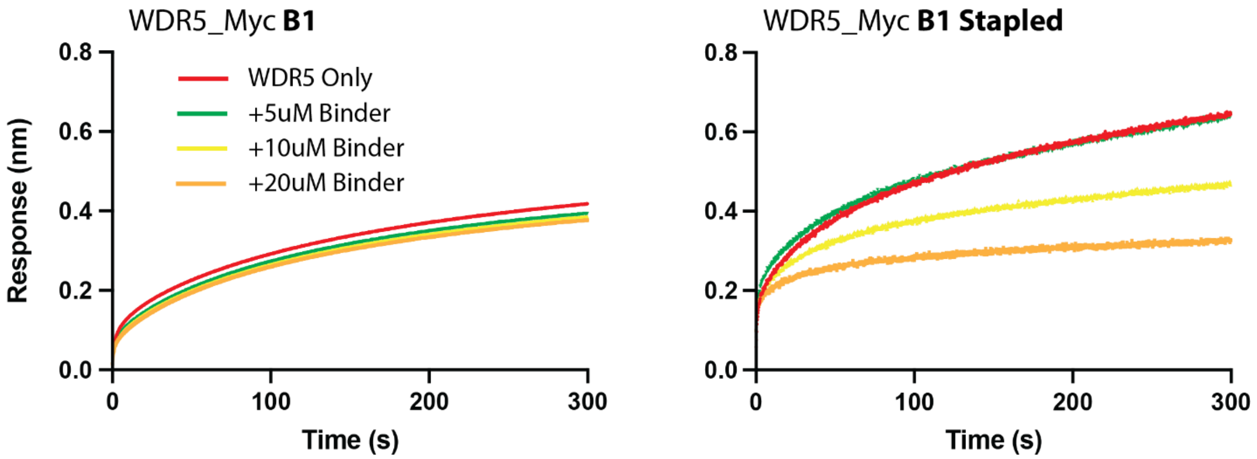


**Supplementary Figure 3: Competition assay for WDR5 with binders B1 and B1 stapled.** Competition analysis (BLI association) of WDR5_Myc Binder1 vs its stapled variant to WDR5-Myc interaction. A Myc-peptide motif was immobilized on BLI tips and was dipped into solutions containing 2.5 μM WDR5 and different concentrations of the binders (5, 10, and 20 μM). Increasing the concentration of the binders in solution causes occupancy of the WDR5-Myc binding pocket and results in a concentration-dependent decrease in BLI response.

**Supplementary Table 4.** Overview of BindCraft input settings used for different protein targets.

| Target Protein | PDB | Target Hotspot | Peptide Length |
|---|---|---|---|
| MDM2 | 1YCR | 73-94 | 10-20 |
| WDR5_WIN | 6DY7 | 133 | 10-20 |
| WDR5_Myc | 6DY7 | 240 | 10-20 |
| PD1 | 4ZQK, chain B | 66 | 10-20 |
| PDL1 | 4ZQK, chain A | 111-127 | 10-20 |

**Supplementary Table 5.** WDR5 Myc B1 stapled sequence.

| Peptide | Native sequence | Cys substitutions |
|---|---|---|
| WDR5_Myc_B1 | DDEDFEQFMKDLDEFLK | DDEDF**C**QFM**C**DLDEFLK |

### Supporting Figure 4: LC-MS Analysis MDM2_B1

**MDM2_B1:**

Ac-K(Biotin)-βA GQSWVNKFMEDMMS-$CONH_2$

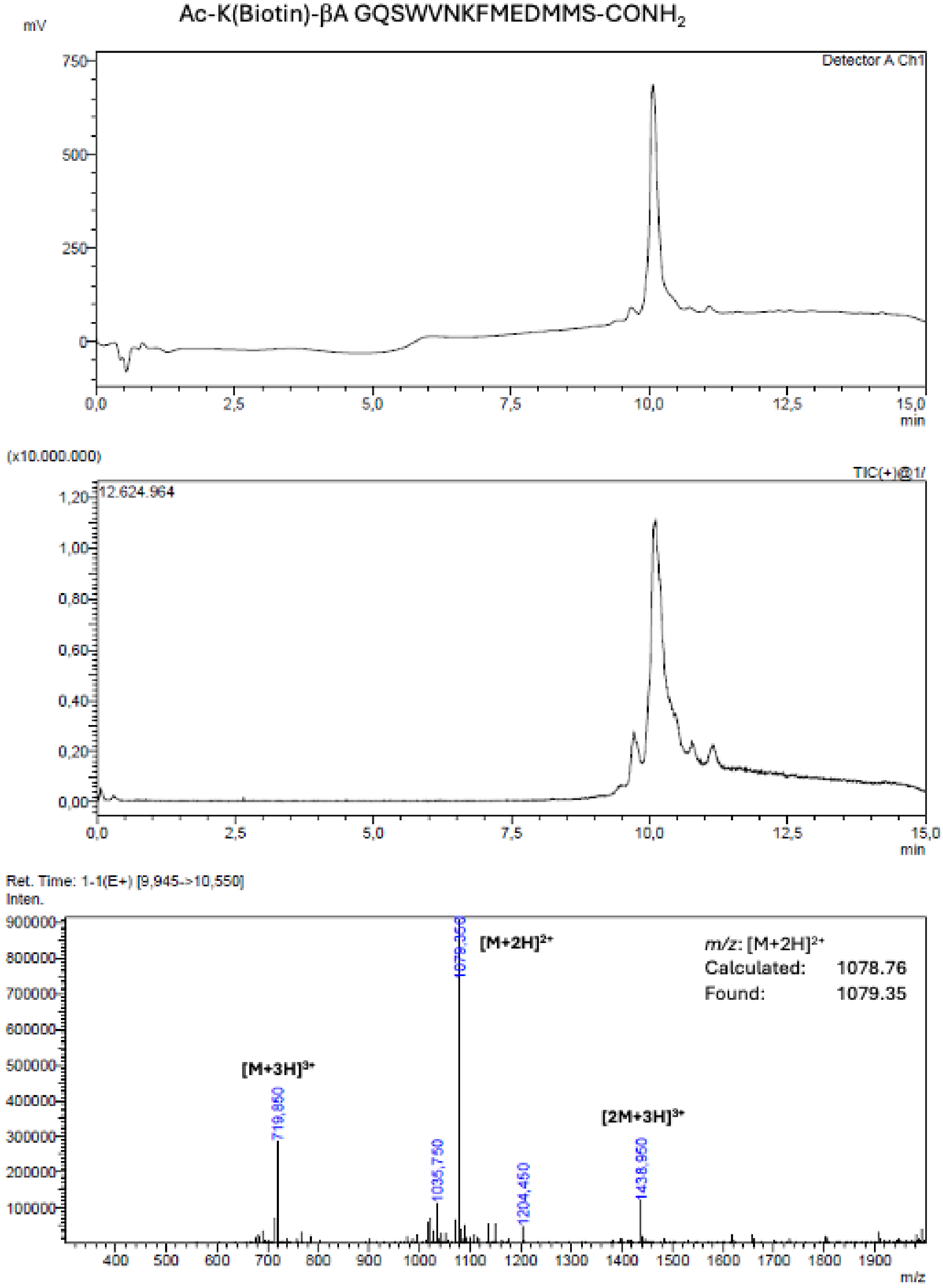

**Supporting Figure 5: LC-MS Analysis MDM2_B2**

**MDM2_B2:**

Ac-K(Biotin)-βA-GQSWVNKFMQDMMS -$CONH_2$

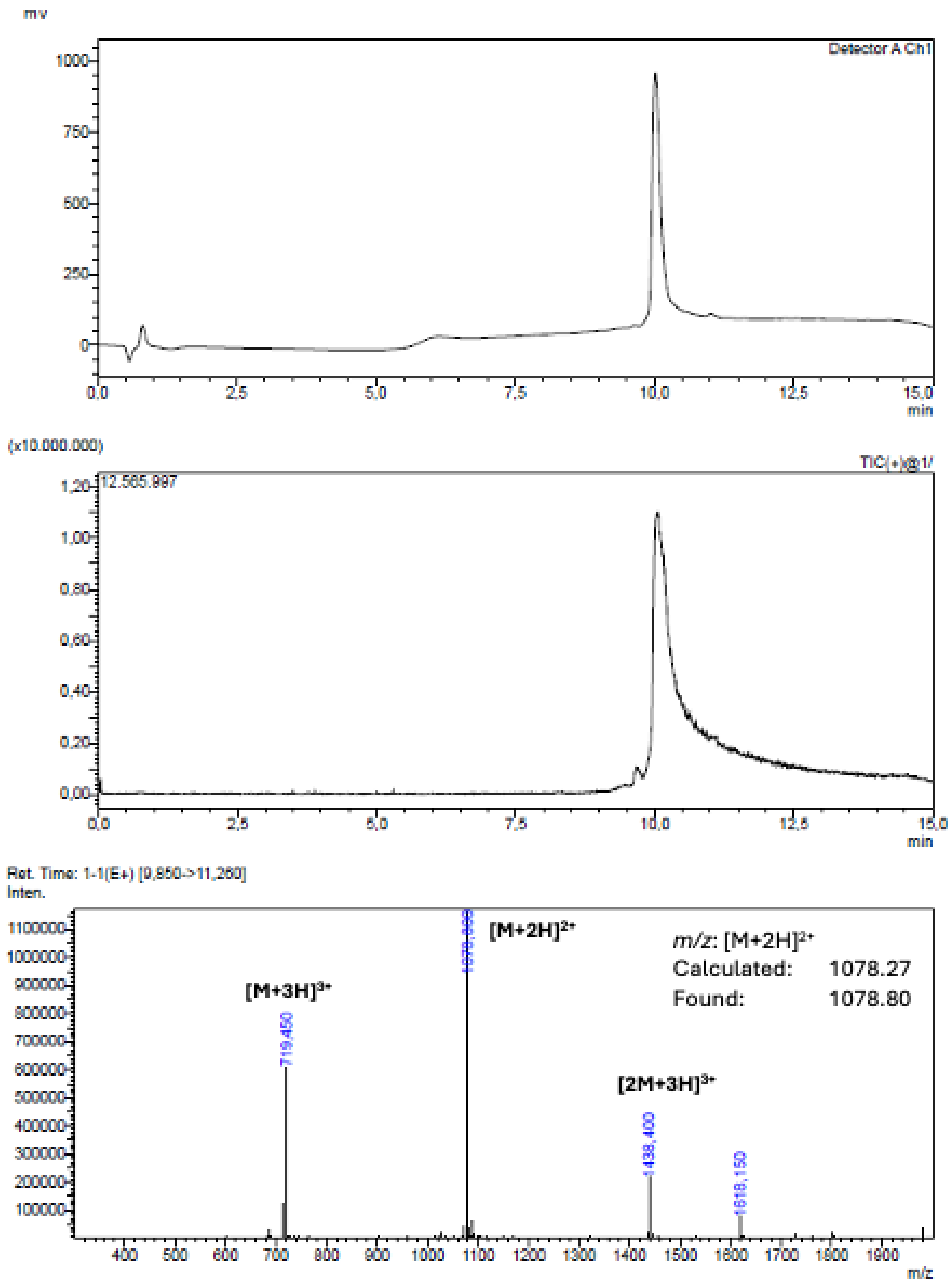

**Supporting Figure 6: LC-MS Analysis MDM2_B3**

**MDM2_B3:**

Ac-K(Biotin)-βA-SDLSGFGKLWEDLMNS -$CONH_2$

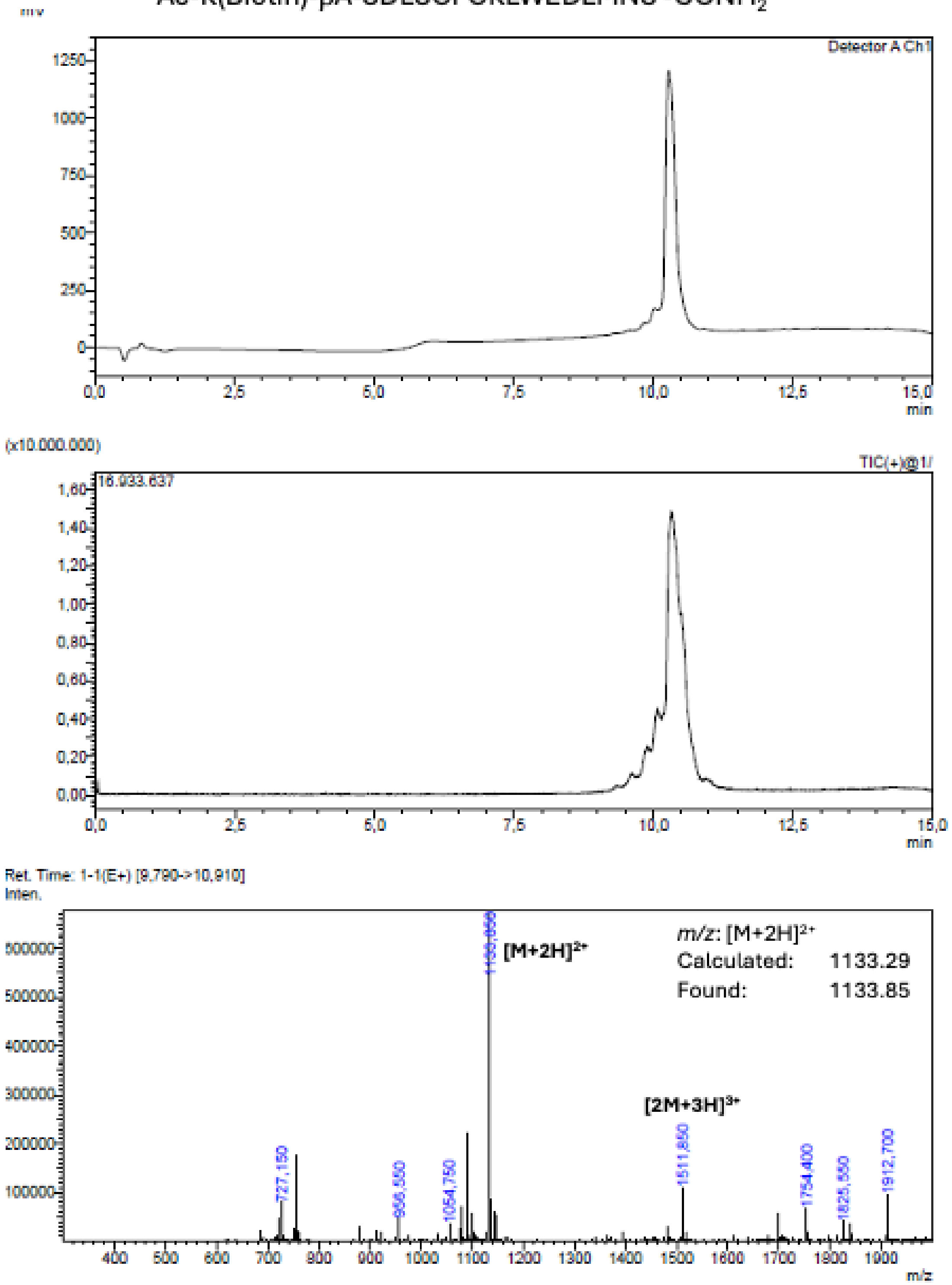

**Supporting Figure 7: LC-MS Analysis MDM2_B4**

**MDM2_B4:**
Ac-K(Biotin)-βA-SDLSGFGKLWQDLMNS -$CONH_2$

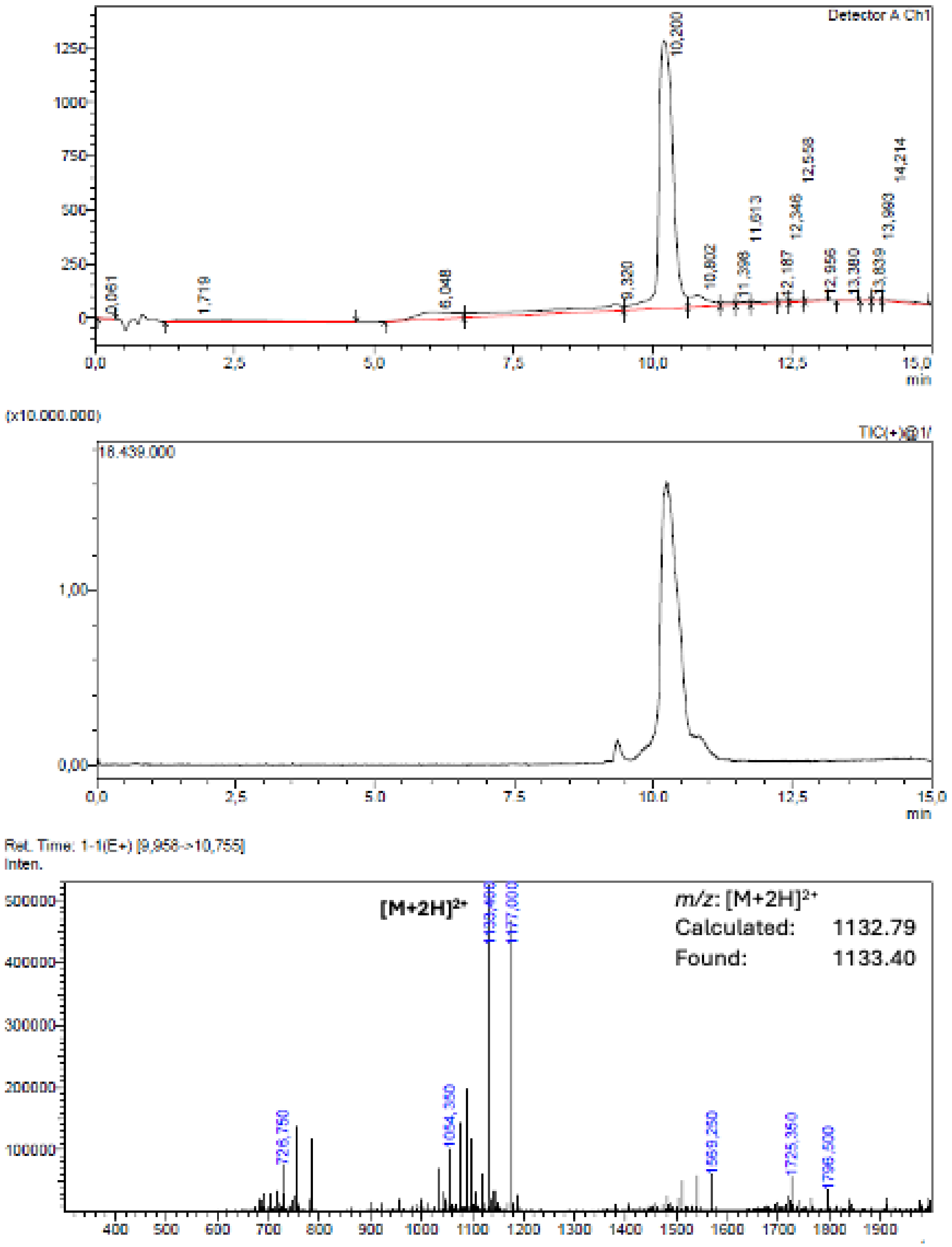

**Supporting Figure 8: LC-MS Analysis MDM2_B5**

**MDM2_B5:**

Ac-K(Biotin)-βA-SASDEFQKEWEDLMNF -$CONH_2$

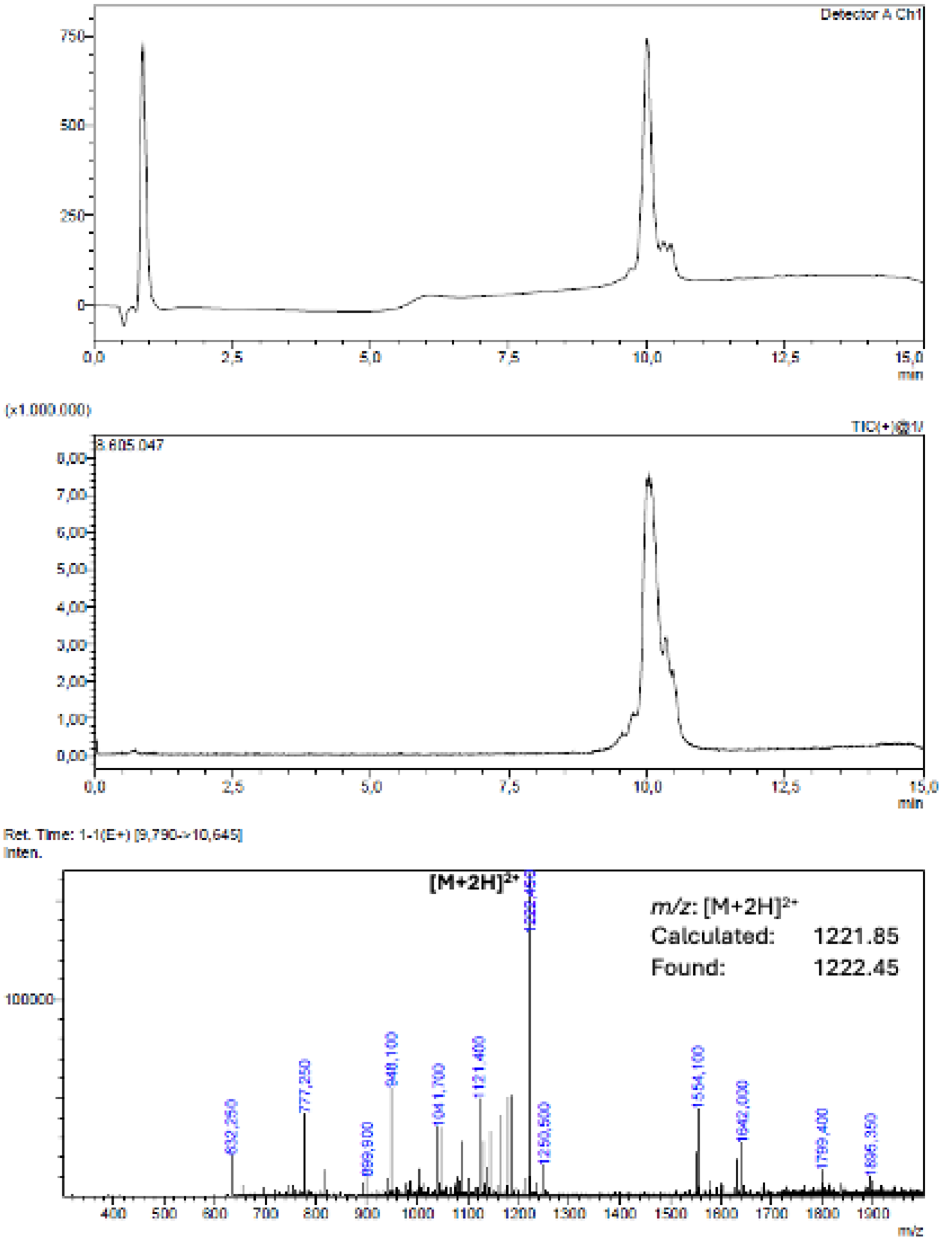

**Supporting Figure 9: LC-MS Analysis MDM2_B6**

**MDM2_B6:**

Ac-K(Biotin)-βA-SASDEFQKEWQDLMNF-$CONH_2$

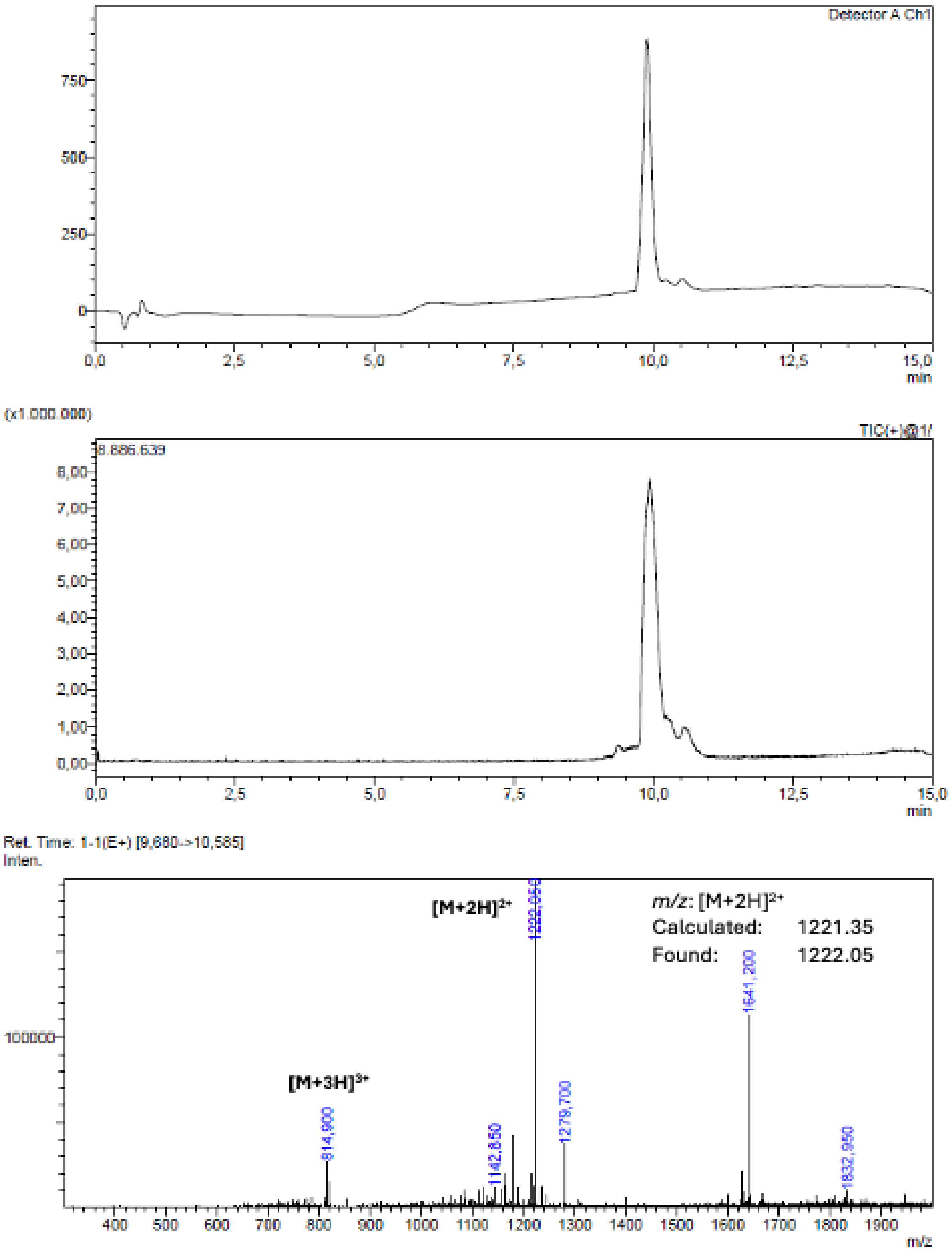

**Supporting Figure 10: LC-MS Analysis MDM2_B8**

**MDM2_B8:**
Ac-K(Biotin)-βA-GTILEWMLNTLEEWRKLS-$CONH_2$

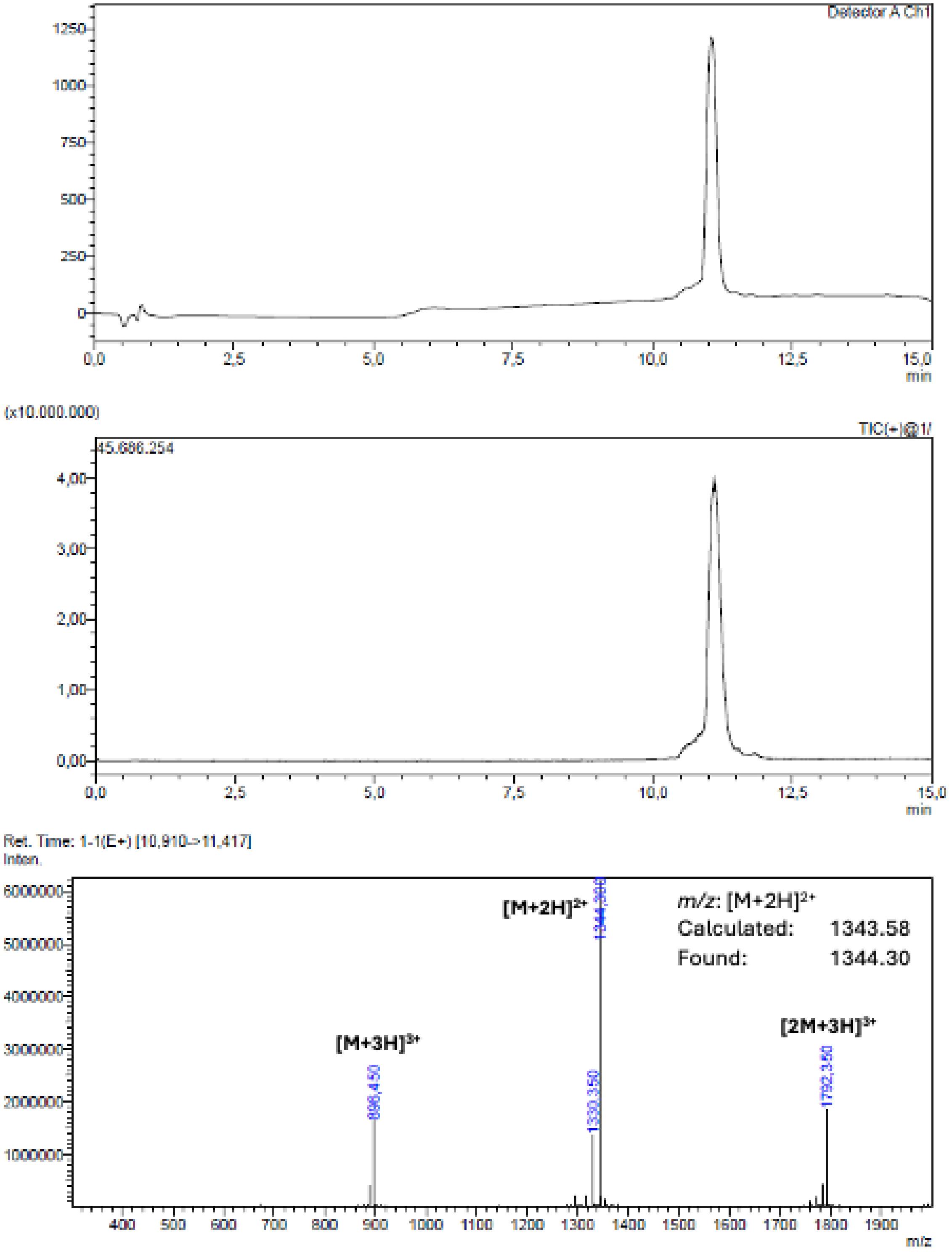

**Supporting Figure 11: LC-MS Analysis MDM2_B11**

**MDM2_B11:**

Ac-K(Biotin)-βA-TFSEQWQELLNS-$CONH_2$

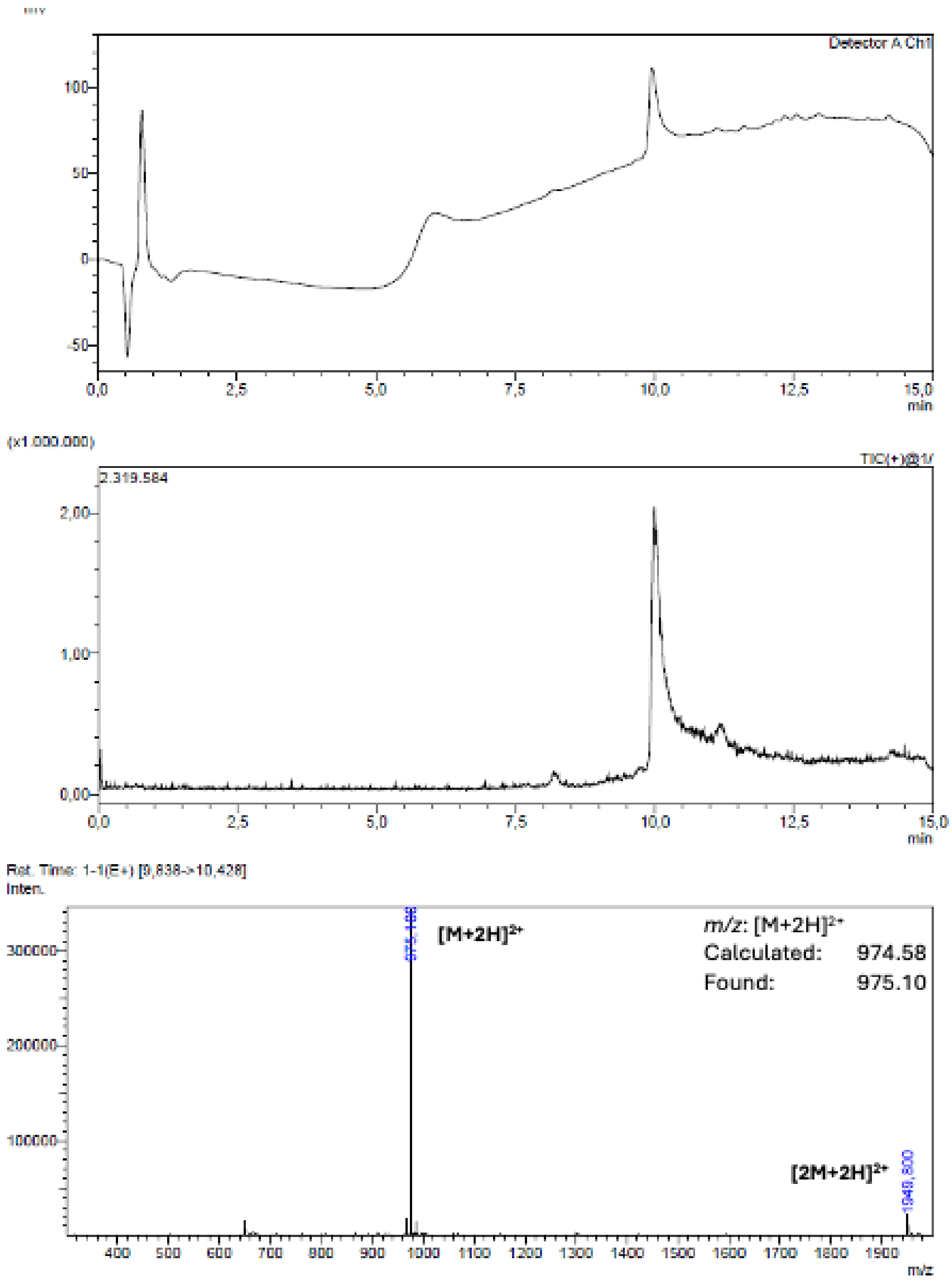

**Supporting Figure 12: LC-MS Analysis MDM2_B12**

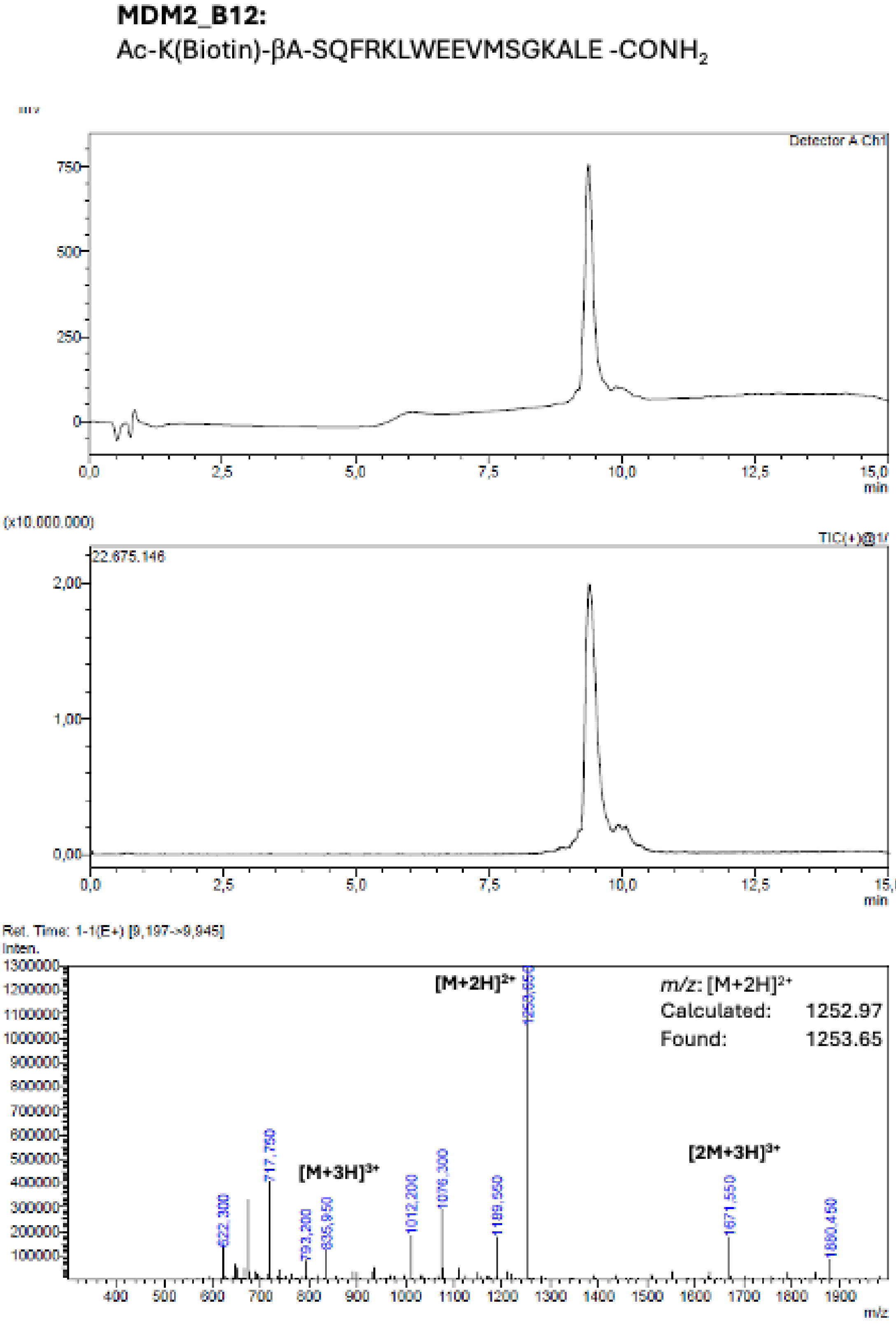

**Supporting Figure 13: LC-MS Analysis MDM2_B13**

**MDM2_B13:**
Ac-K(Biotin)-βA-SQFRELWEEVMSGKALE-$CONH_2$

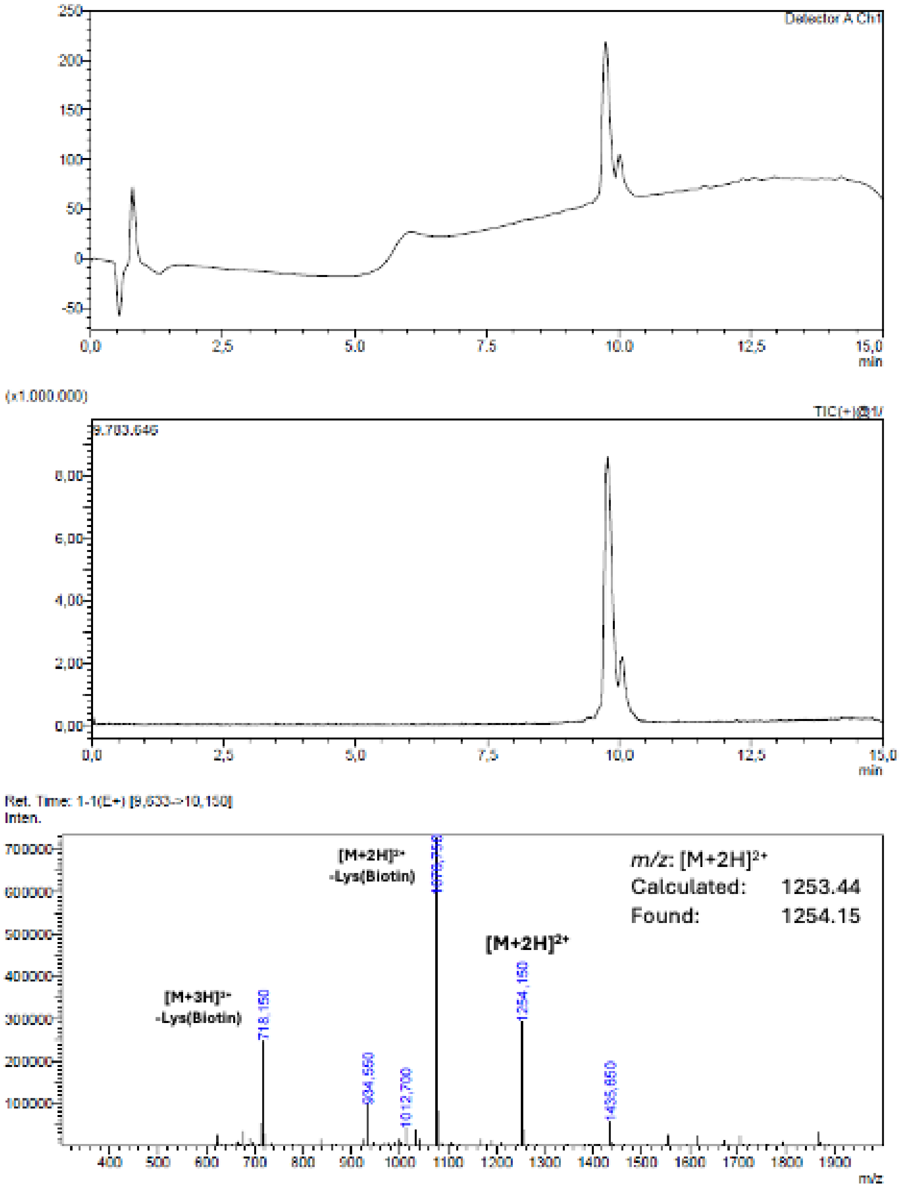

**Supporting Figure 14: LC-MS Analysis MDM2_B14**

**MDM2_B14:**
Ac-K(Biotin)-βA-TGFQKEWEKVMNS-$CONH_2$

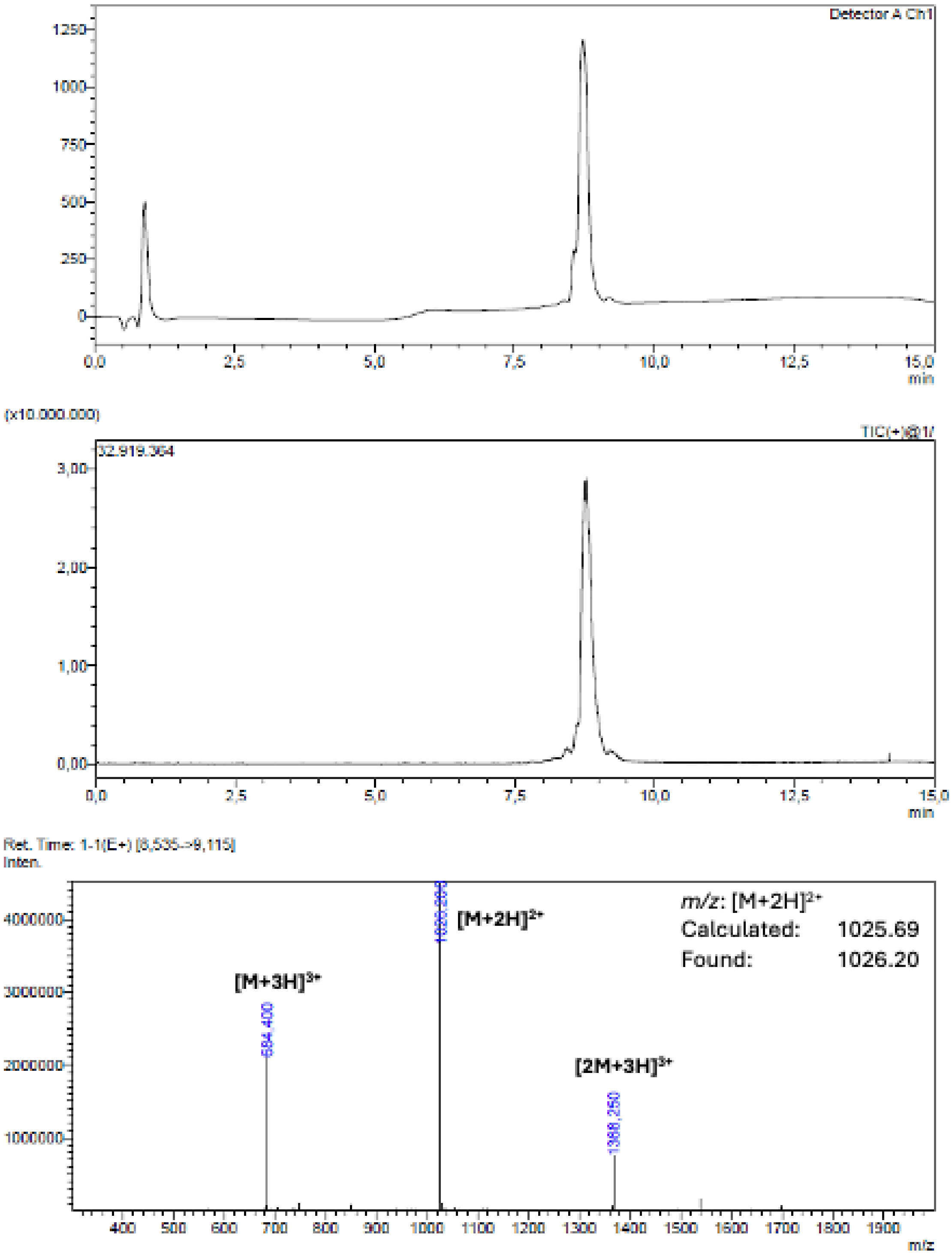

**Supporting Figure 15: LC-MS Analysis MDM2_B15**

**MDM2_B15:**
Ac-K(Biotin)-βA-SPSEFQKHWQDLWNDYMK-$CONH_2$

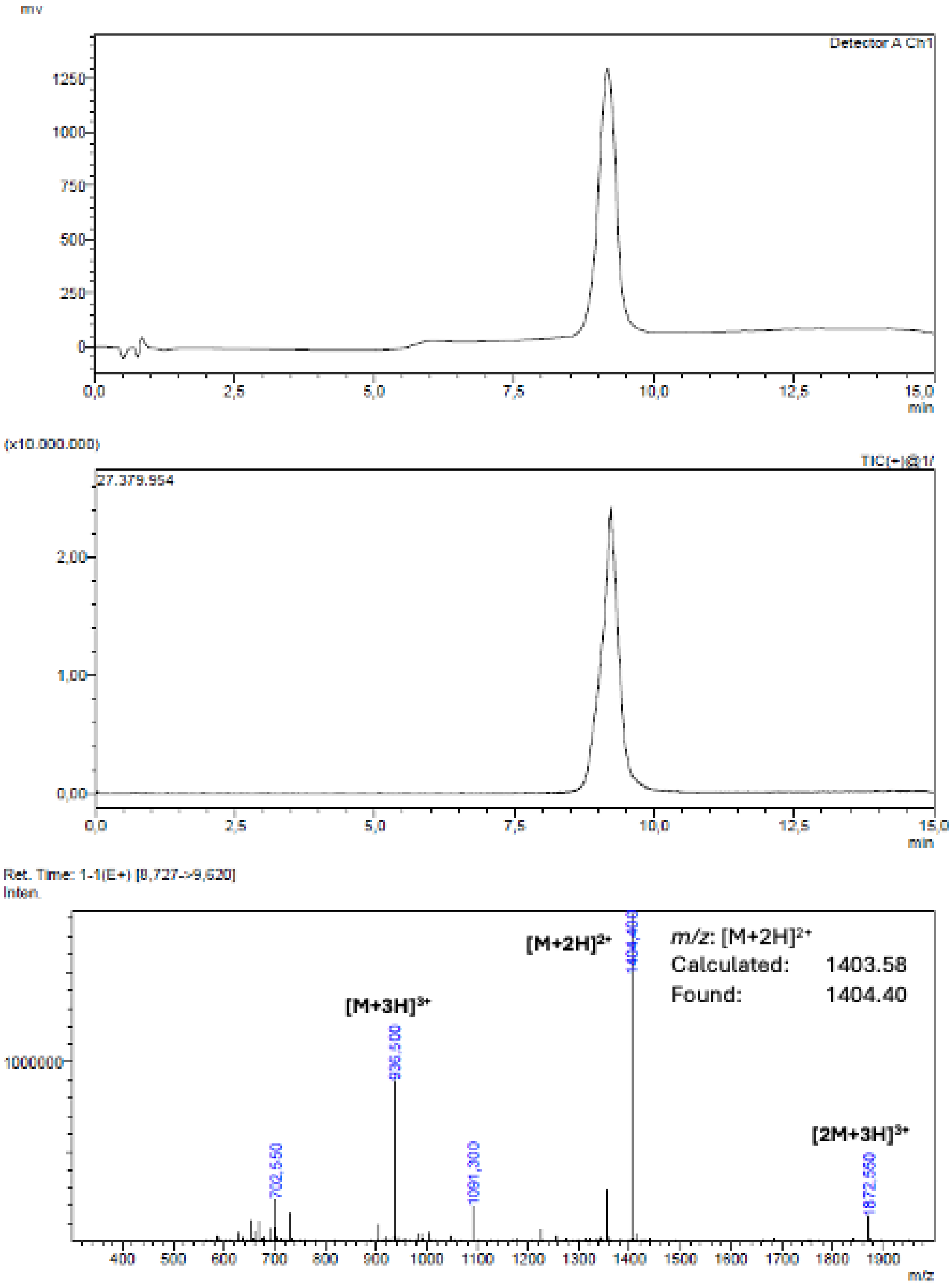

**Supporting Figure 16: LC-MS Analysis MDM2_B17**

**MDM2_B17:**

Ac-K(Biotin)-βA-SPSEFQKHWQDLWDDYMK-$CONH_2$

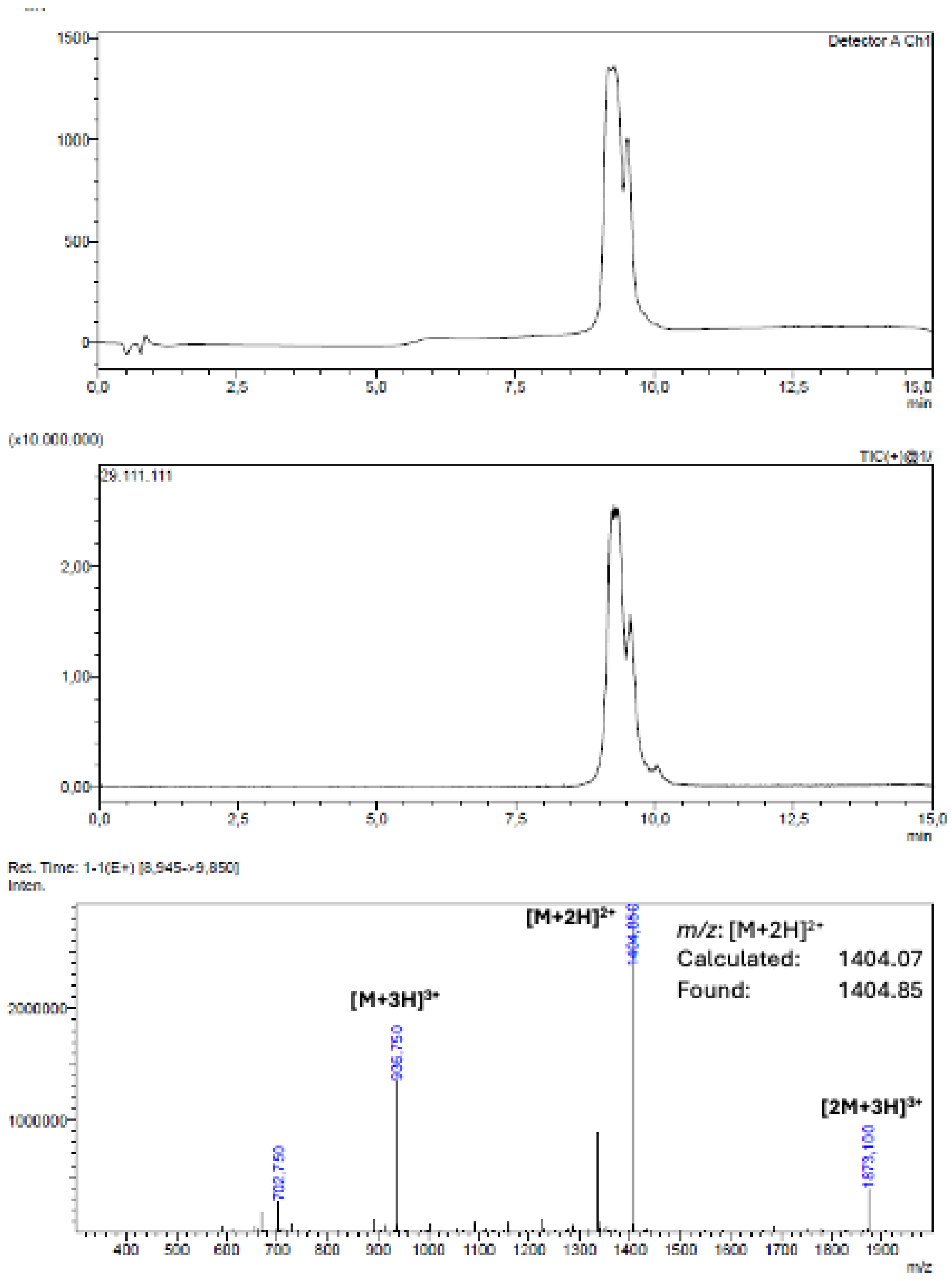

**Supporting Figure 17: LC-MS Analysis MDM2_B18**

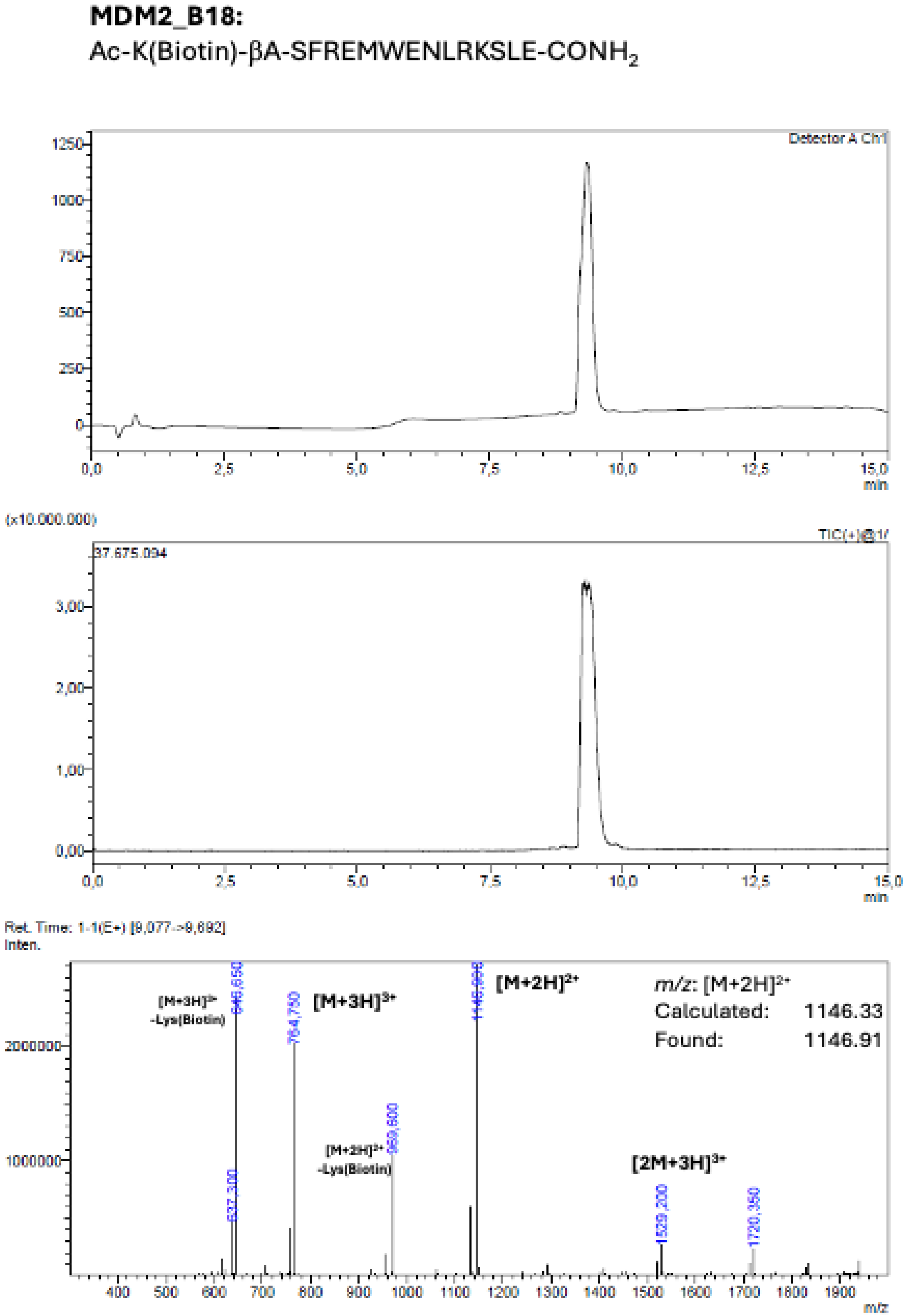

**Supporting Figure 18: LC-MS Analysis MDM2_B19**

**MDM2_B19:**
Ac-K(Biotin)-βA-SFRDMWENLRKSLE-$CONH_2$

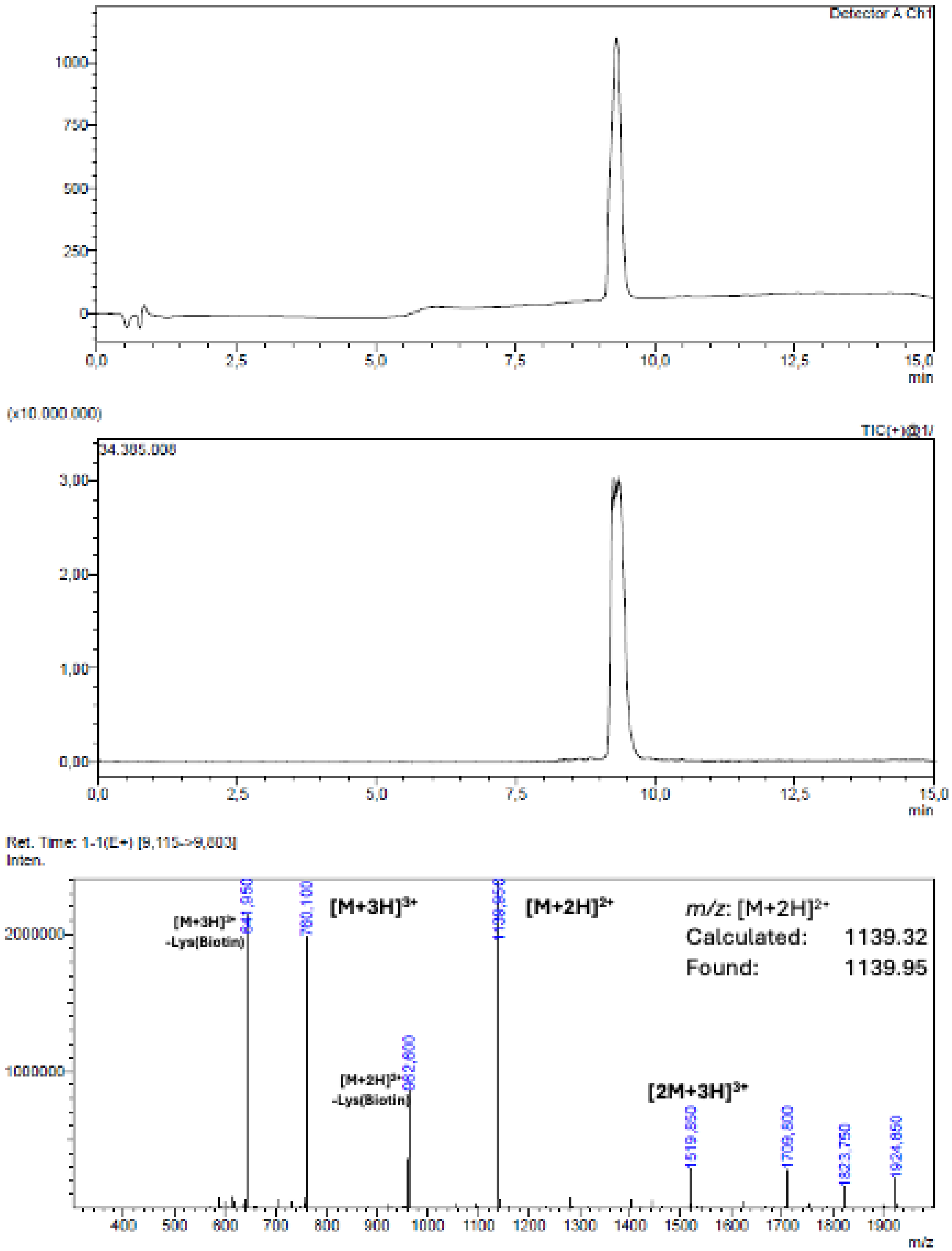

**Supporting Figure 19: LC-MS Analysis MDM2_B20**

**MDM2_B20:**

Ac-K(Biotin)-βA-GWEGFMKQWKEFSENLEKYM-$CONH_2$

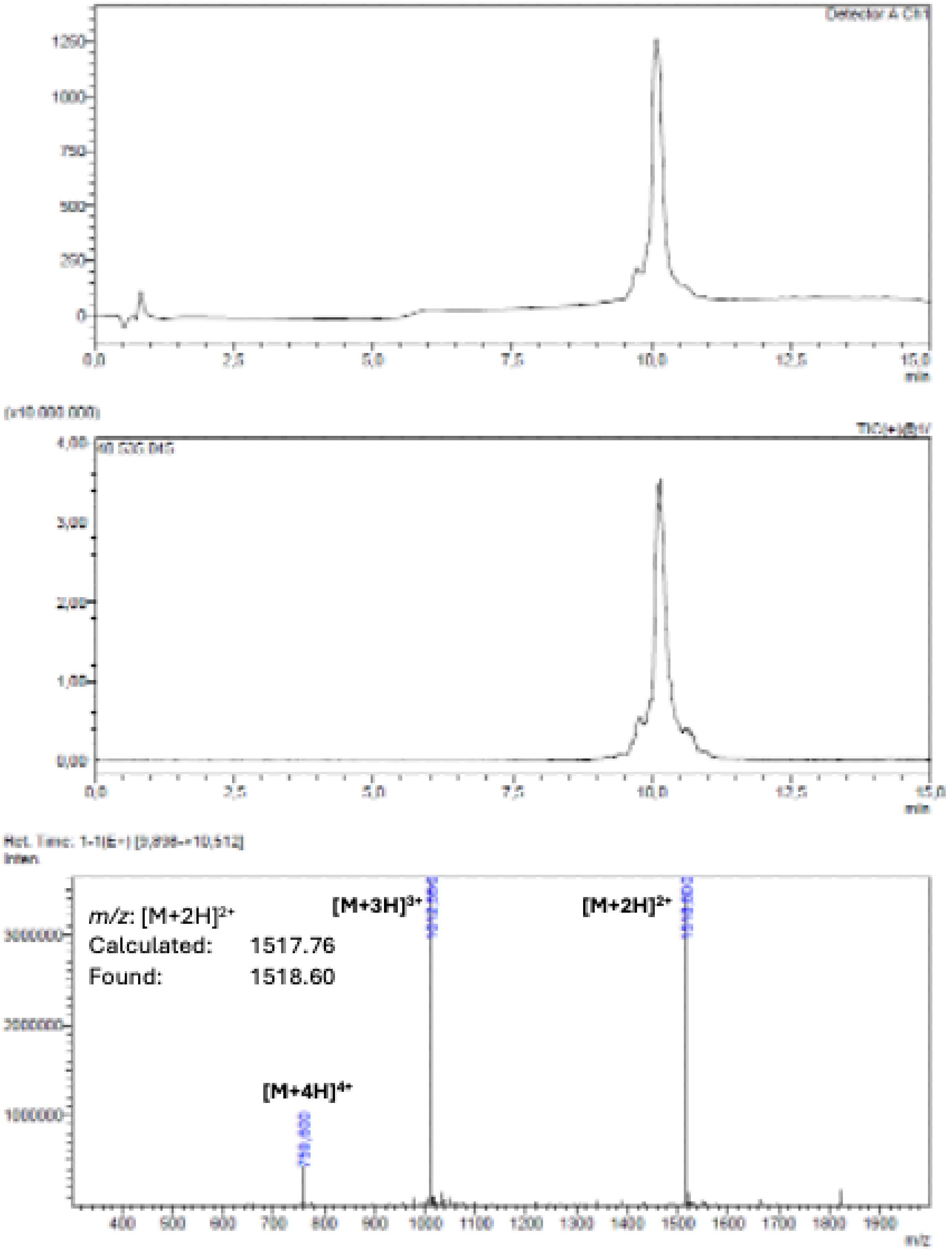

### Supporting Figure 20: LC-MS Analysis WDR5_MYC_B1

**WDR5_Myc_B1:**
Ac-K(Biotin)-βA-DDEDFEQFMKDLDEFLK-$CONH_2$

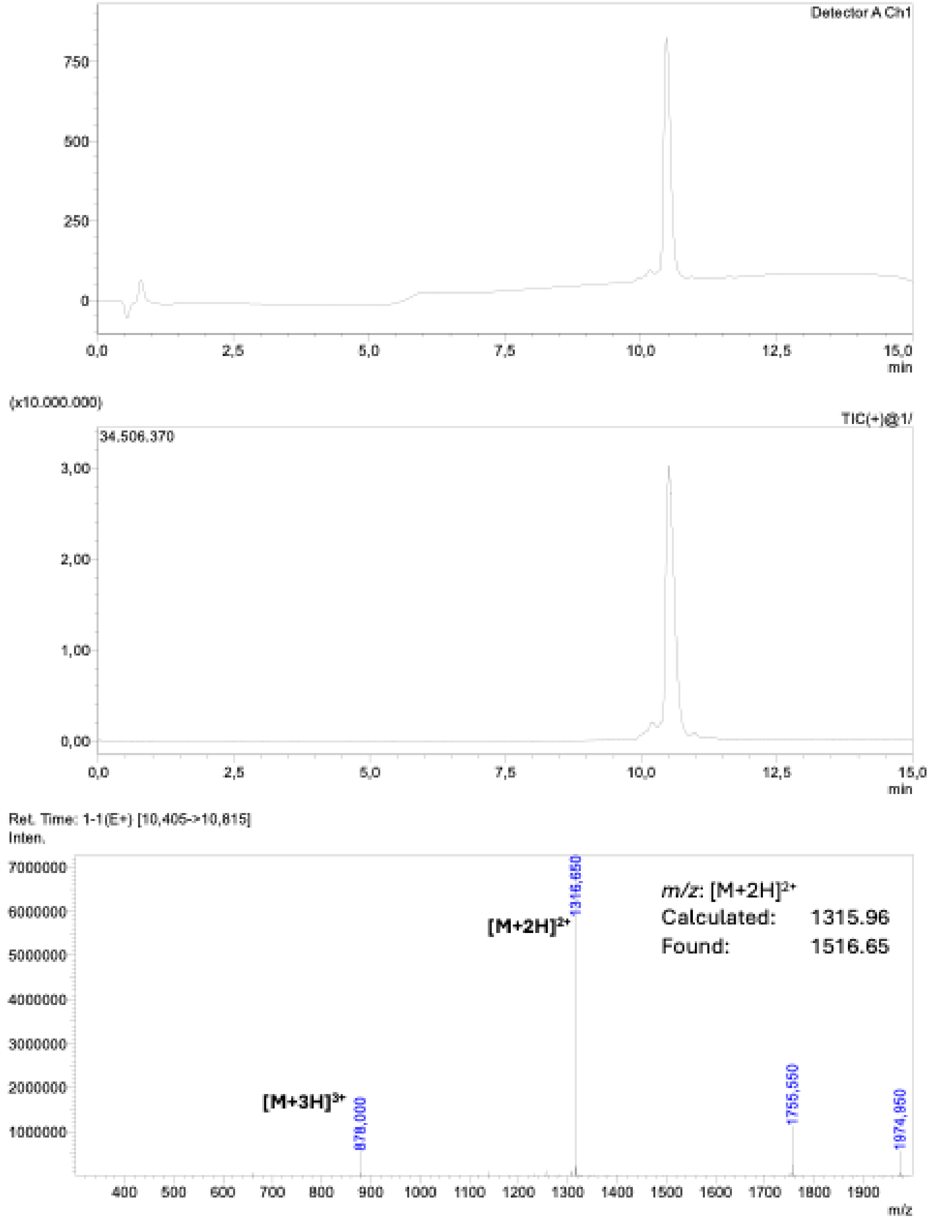

**Supporting Figure 21: LC-MS Analysis WDR5_MYC_B3**

**WDR5_Myc_B3:**
Ac-K(Biotin)-βA-DTDKVLEQIEKEQ-$CONH_2$

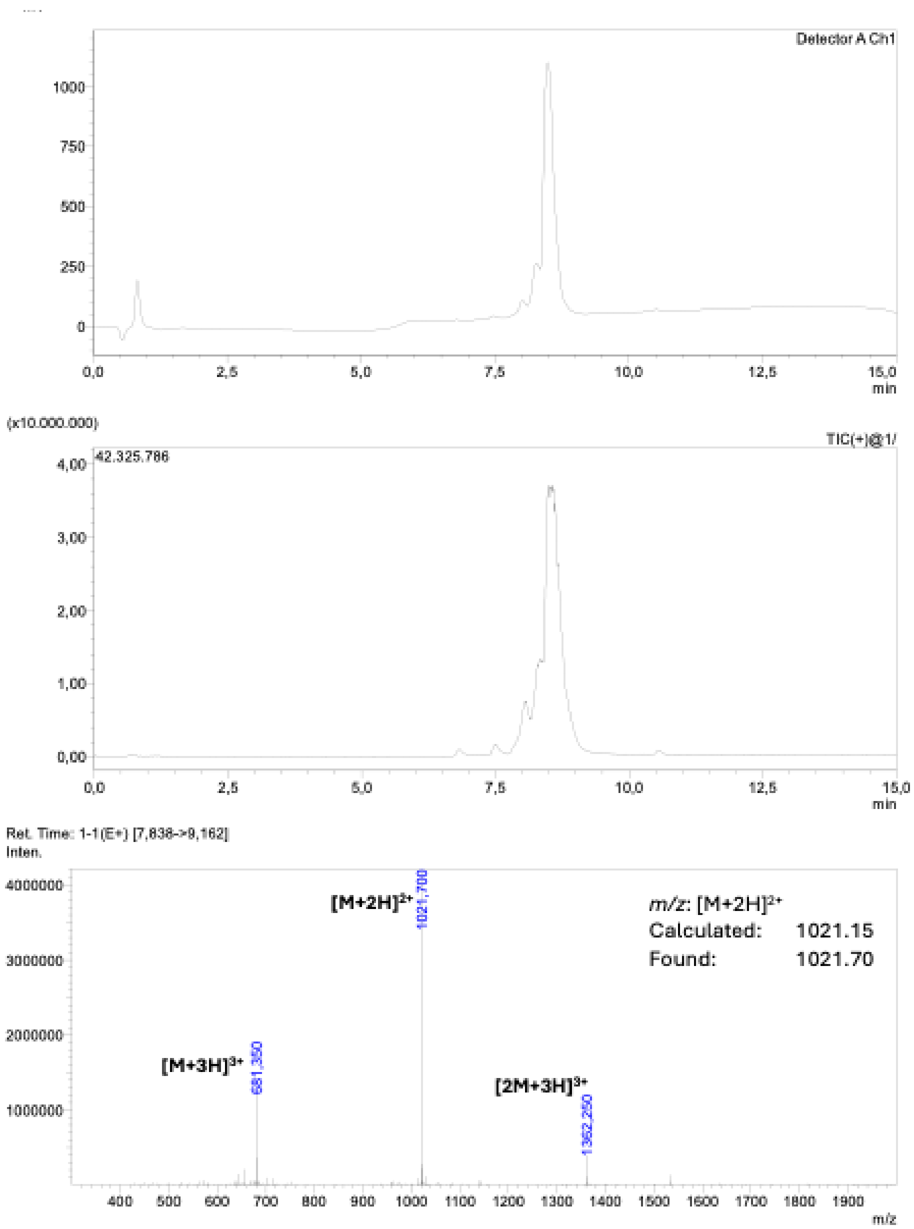

**Supporting Figure 22: LC-MS Analysis WDR5_MYC_B5**

**WDR5_Myc_B5:**

Ac-K(Biotin)-βA-NPYQEIYDKQAAEFDKFMS-$CONH_2$

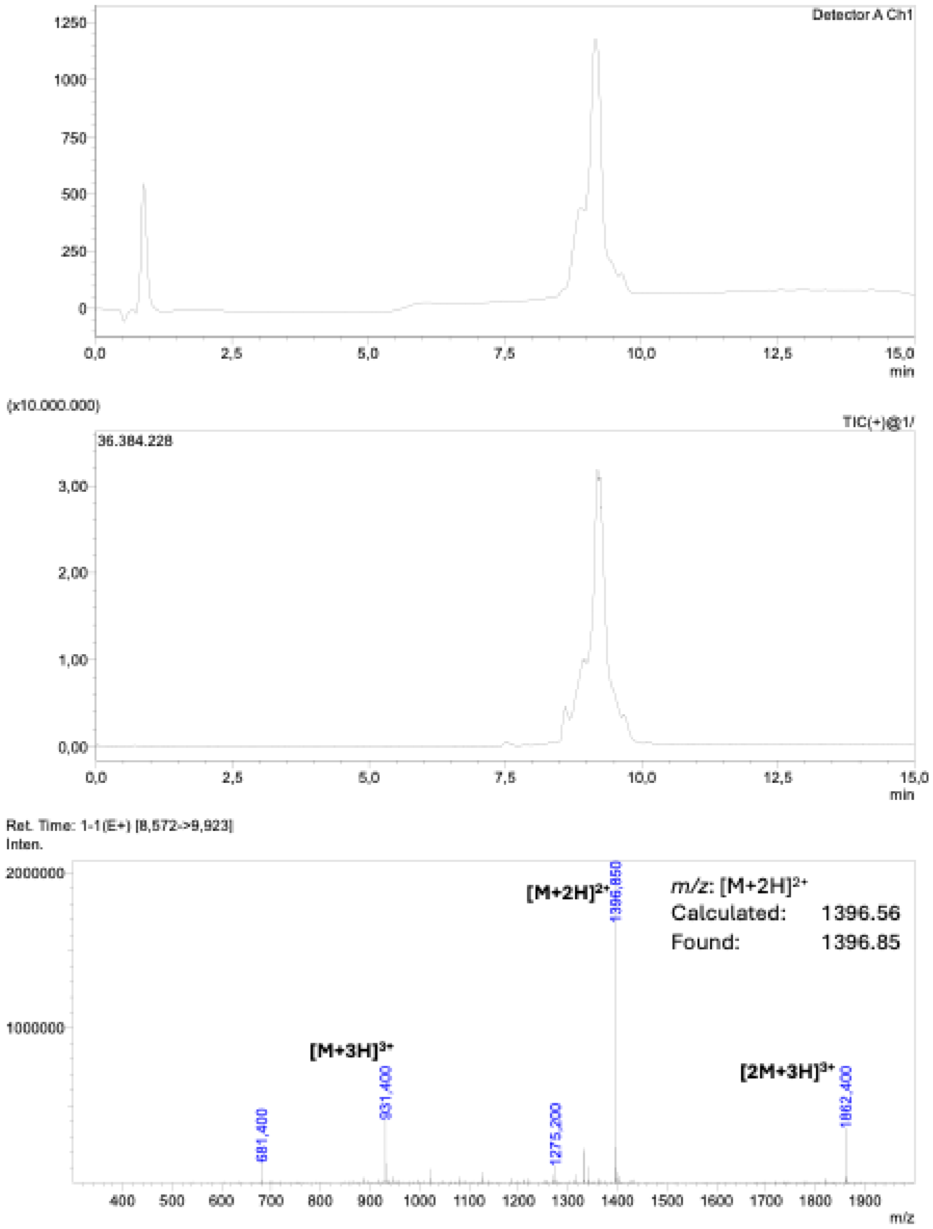

### Supporting Figure 23: LC-MS Analysis WDR5_MYC_B6

**WDR5_Myc_B6:**
Ac-K(Biotin)-βA-SAEEDEKAMQELMDYLNSM-$CONH_2$

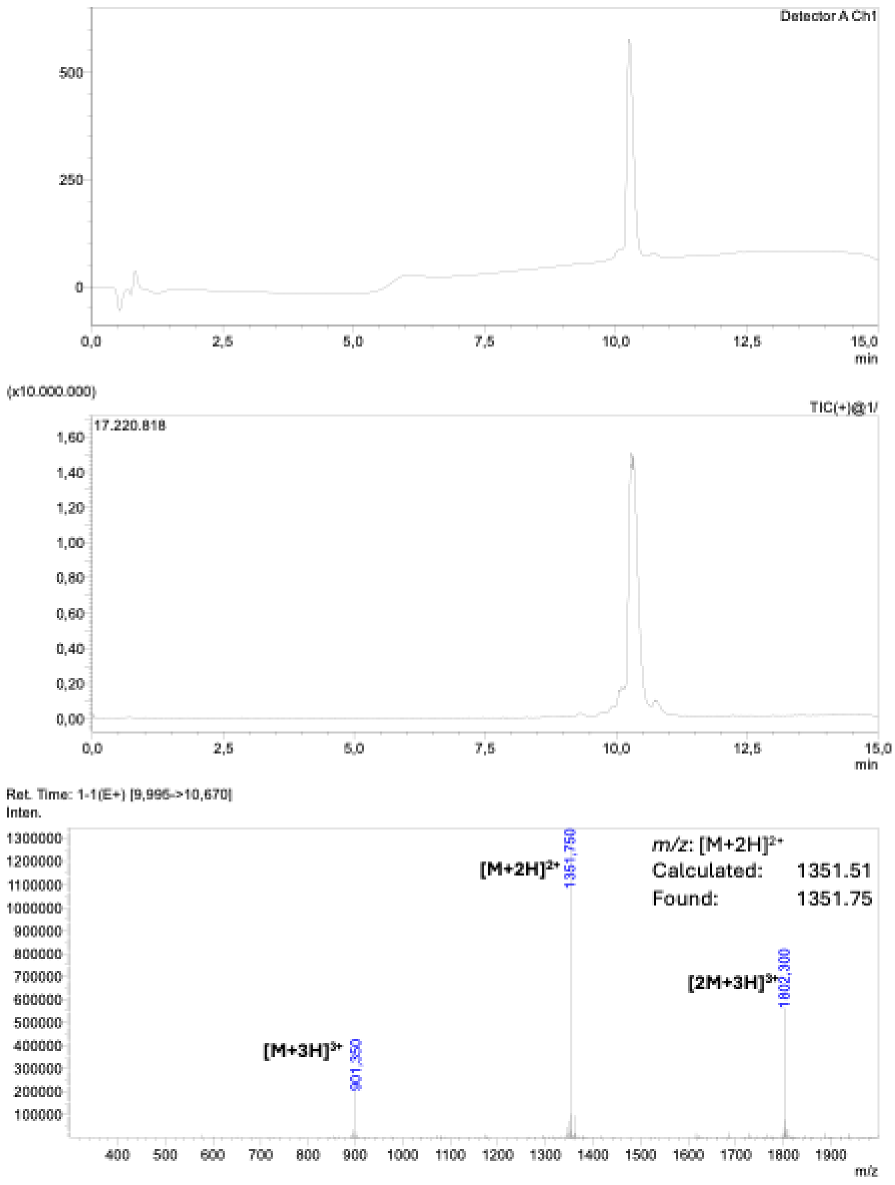

**Supporting Figure 24: LC-MS Analysis WDR5_MYC_B7**

**WDR5_Myc_B7:**
Ac-K(Biotin)-βA-DPEVDELIKFMDENADAWR-$CONH_2$

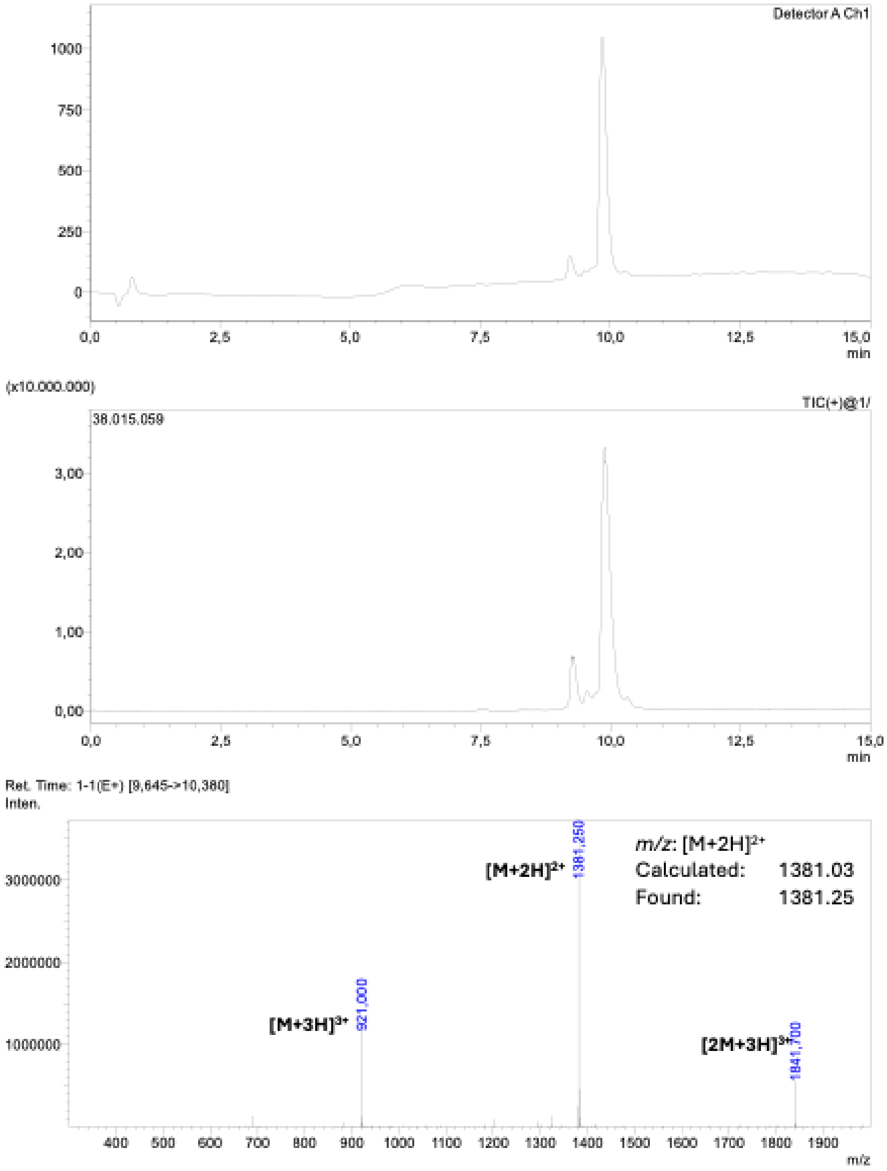

**Supporting Figure 25: LC-MS Analysis WDR5_MYC_B8**

**WDR5_Myc_B8:**
Ac-K(Biotin)-βA-MSPEEDLEEWIRNL-$CONH_2$

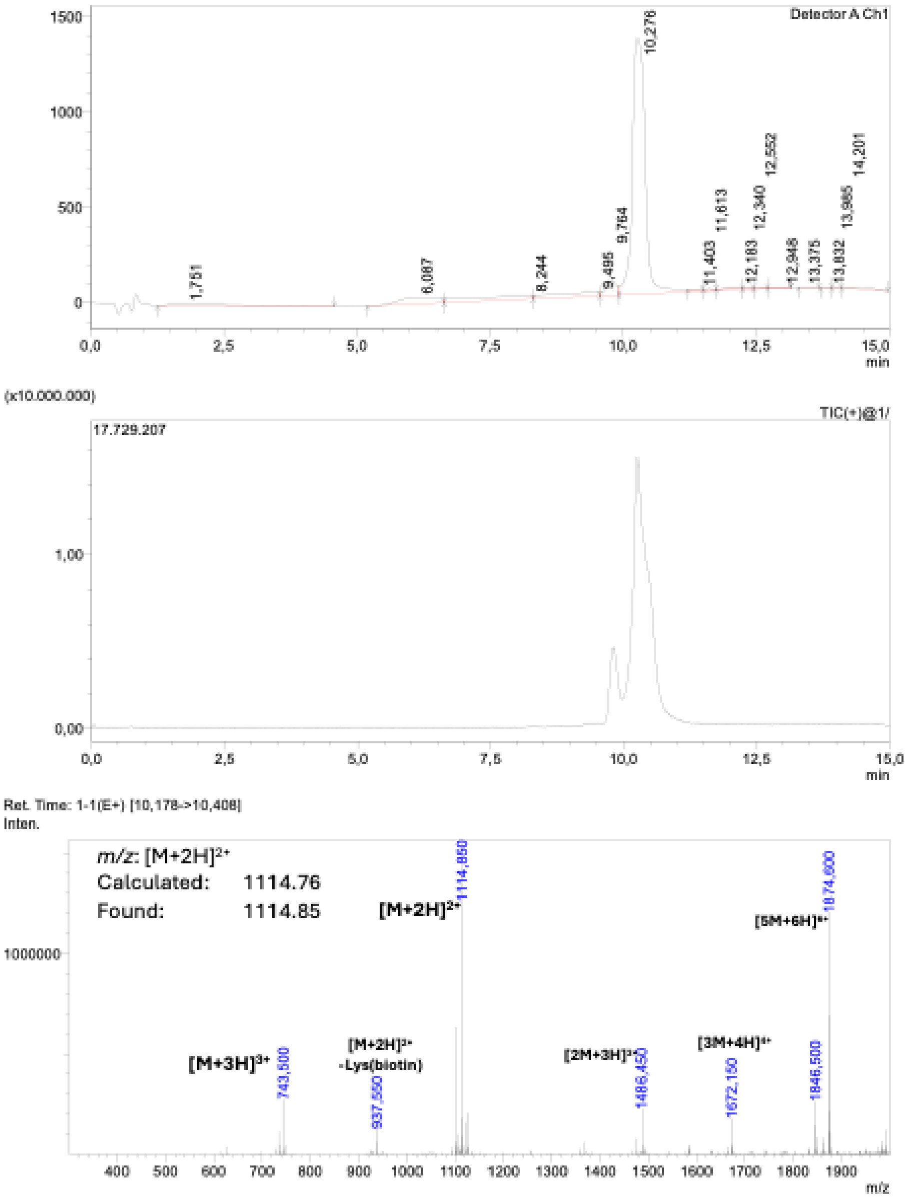

### Supporting Figure 26: LC-MS Analysis WDR5_MYC_B10

**WDR5_Myc_B10:**
Ac-K(Biotin)-βA-SPDKDVQELIDYLNS-$CONH_2$

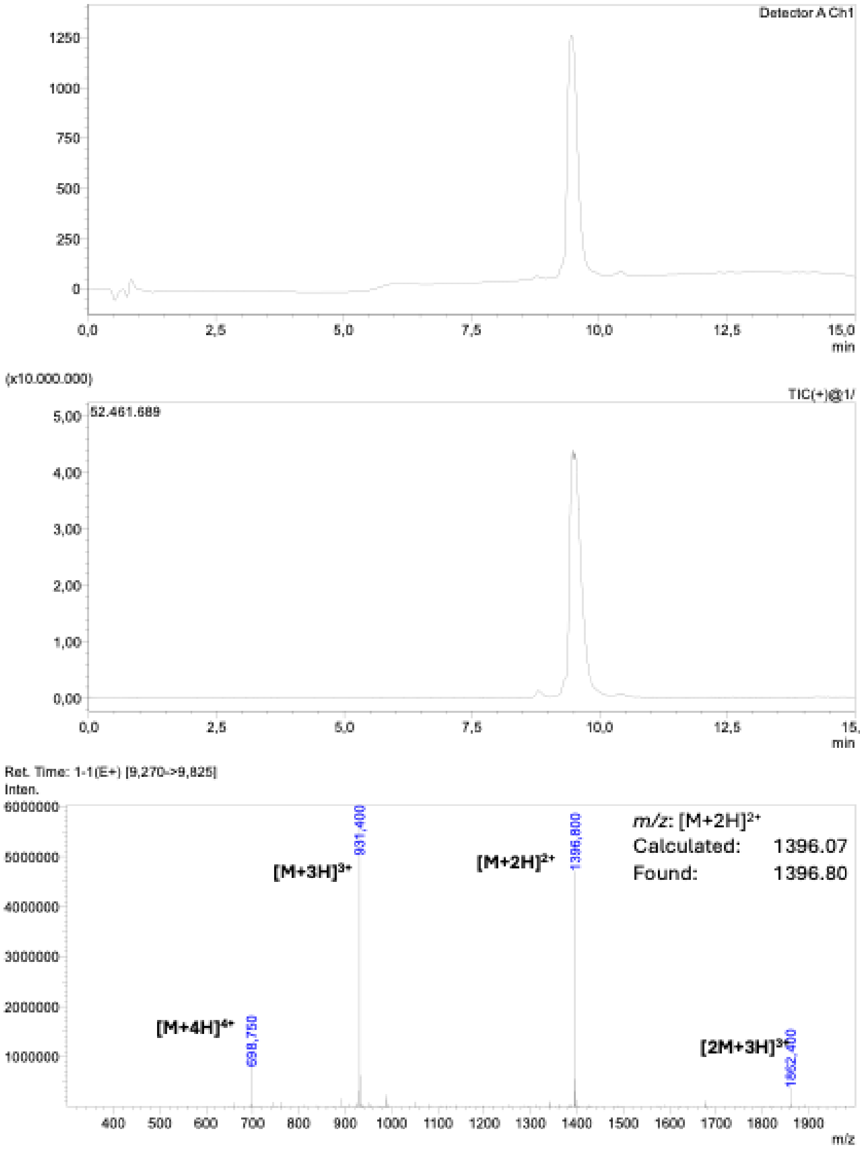

**Supporting Figure 27: LC-MS Analysis WDR5_MYC_B11**

**WDR5_Myc_B11:**
Ac-K(Biotin)-βA-SPDKDVQELIDYLNS-$CONH_2$

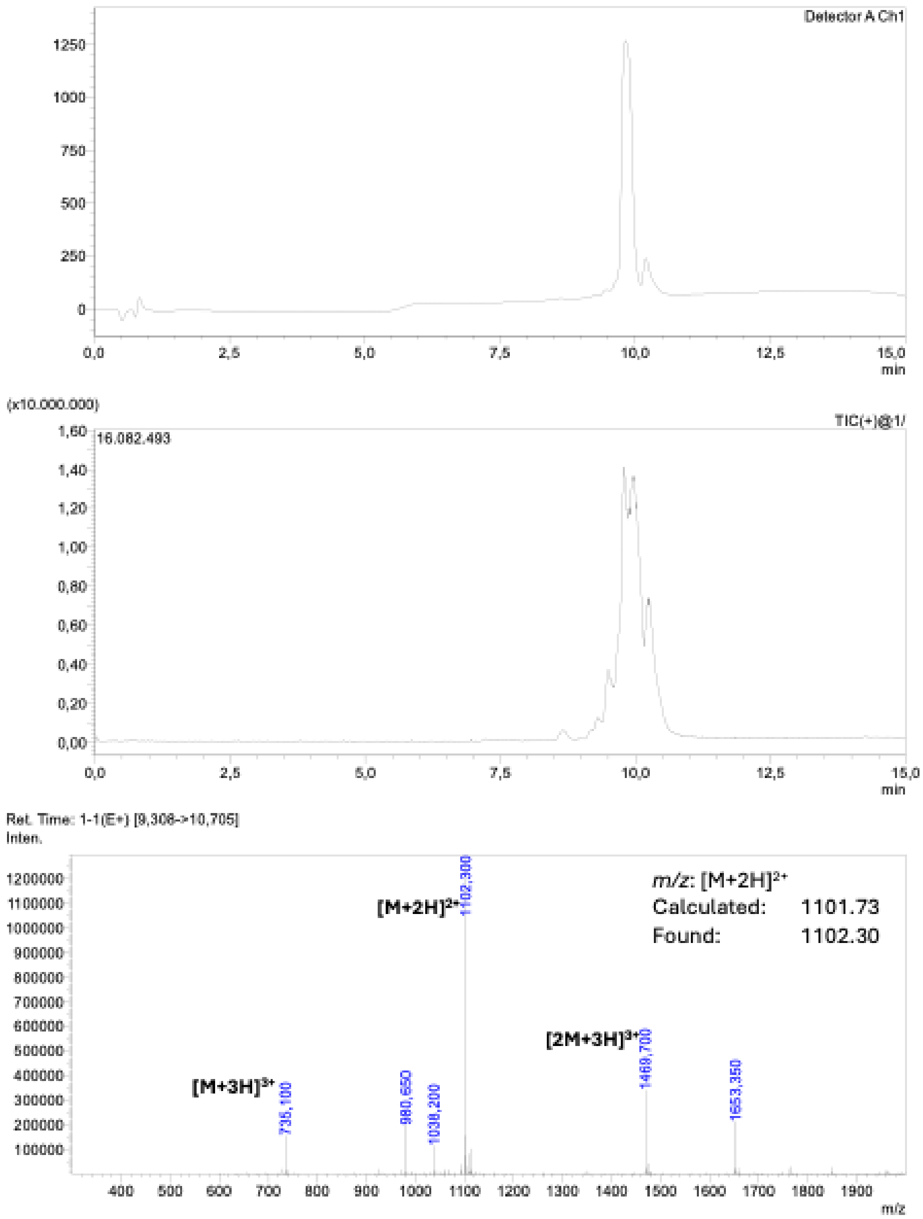

**Supporting Figure 28: LC-MS Analysis WDR5_MYC_B12**

**WDR5_Myc_B12:**
Ac-K(Biotin)-βA-DTDEVLEQIEKEQ-$CONH_2$

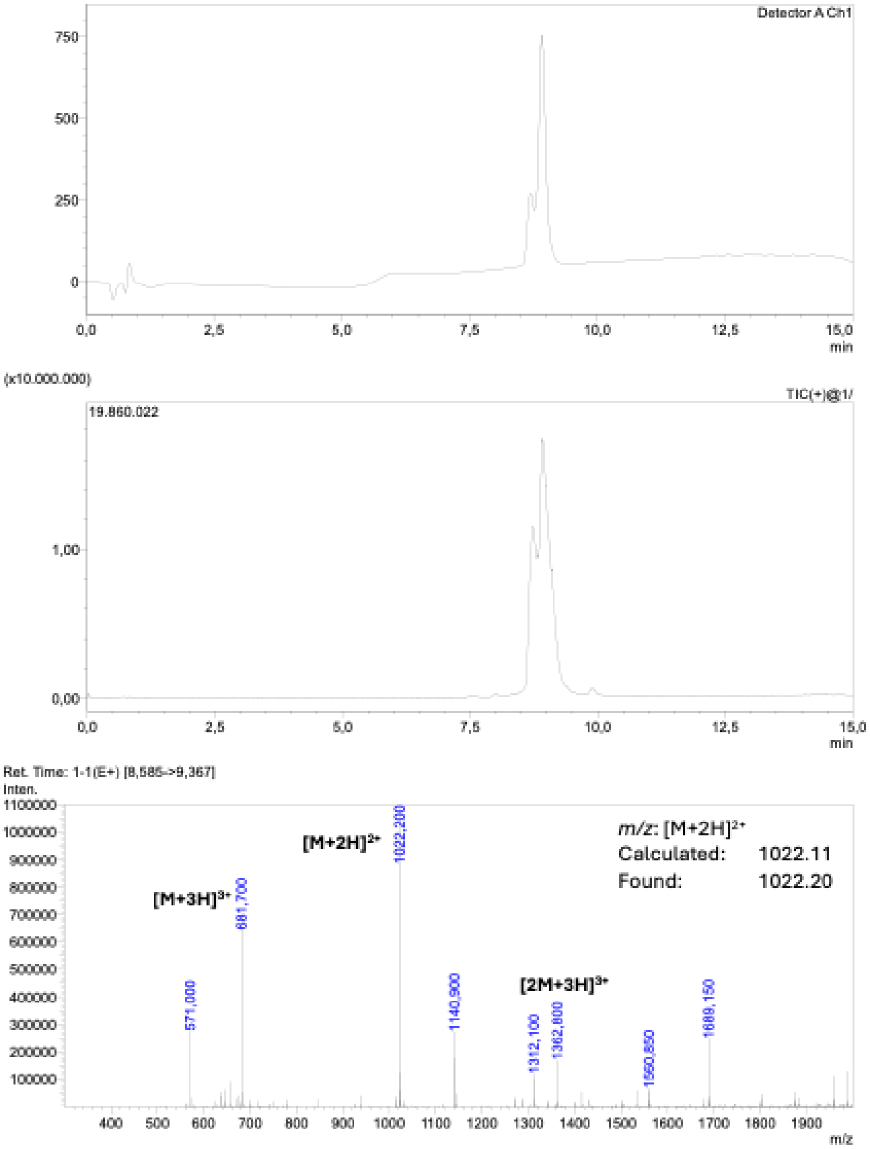

**Supporting Figure 29: LC-MS Analysis WDR5_WIN_B2**

**WDR5_Win_B2:**
Ac-K(Biotin)-βA-SENTKKRFEDMMNDMYK-$CONH_2$

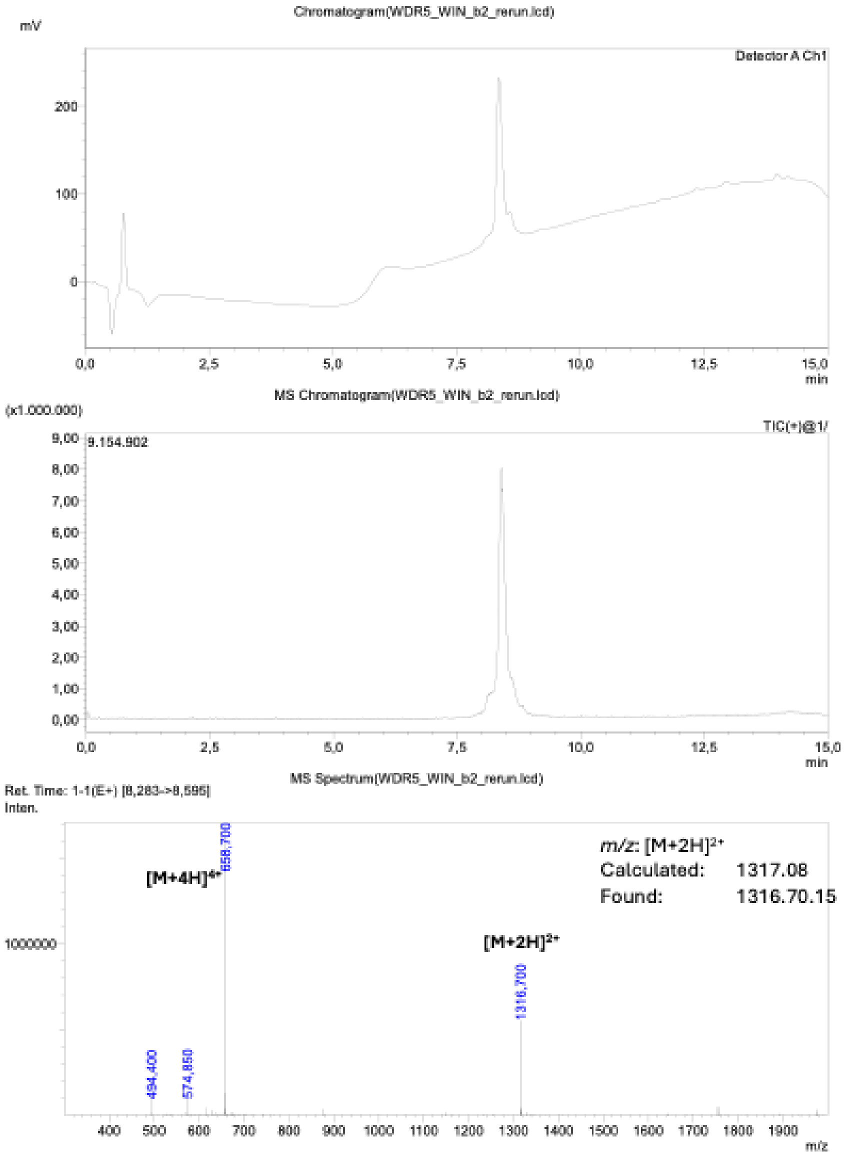

**Supporting Figure 30: LC-MS Analysis WDR5_WIN_B9**

**WDR5_Win_B9:**
Ac-K(Biotin)-βA-MSRGKEQFDKTHNEIHQ-$CONH_2$

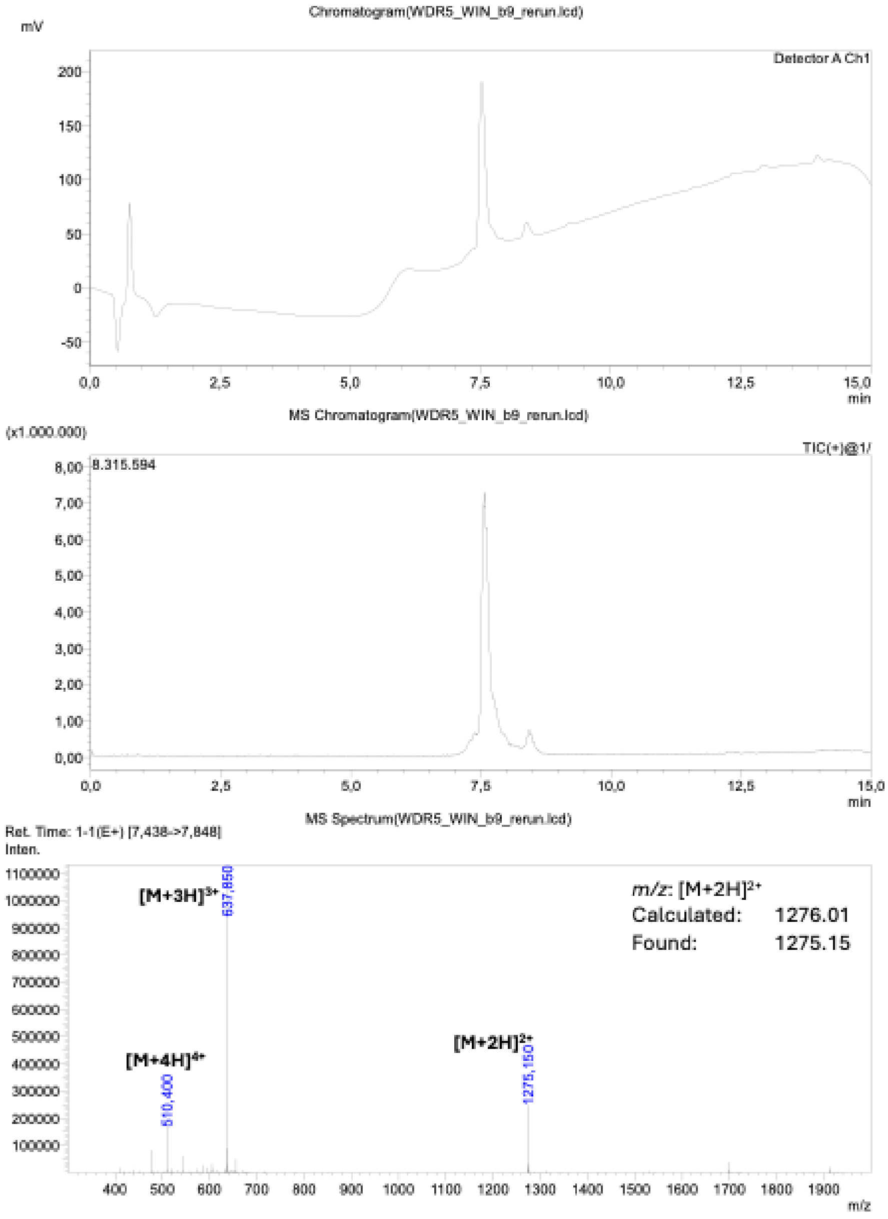

**Supporting Figure 31: LC-MS Analysis WDR5_WIN_B10**

**WDR5_Win_B10:**
Ac-K(Biotin)-βA-DPRKMFEEHRNALW-$CONH_2$

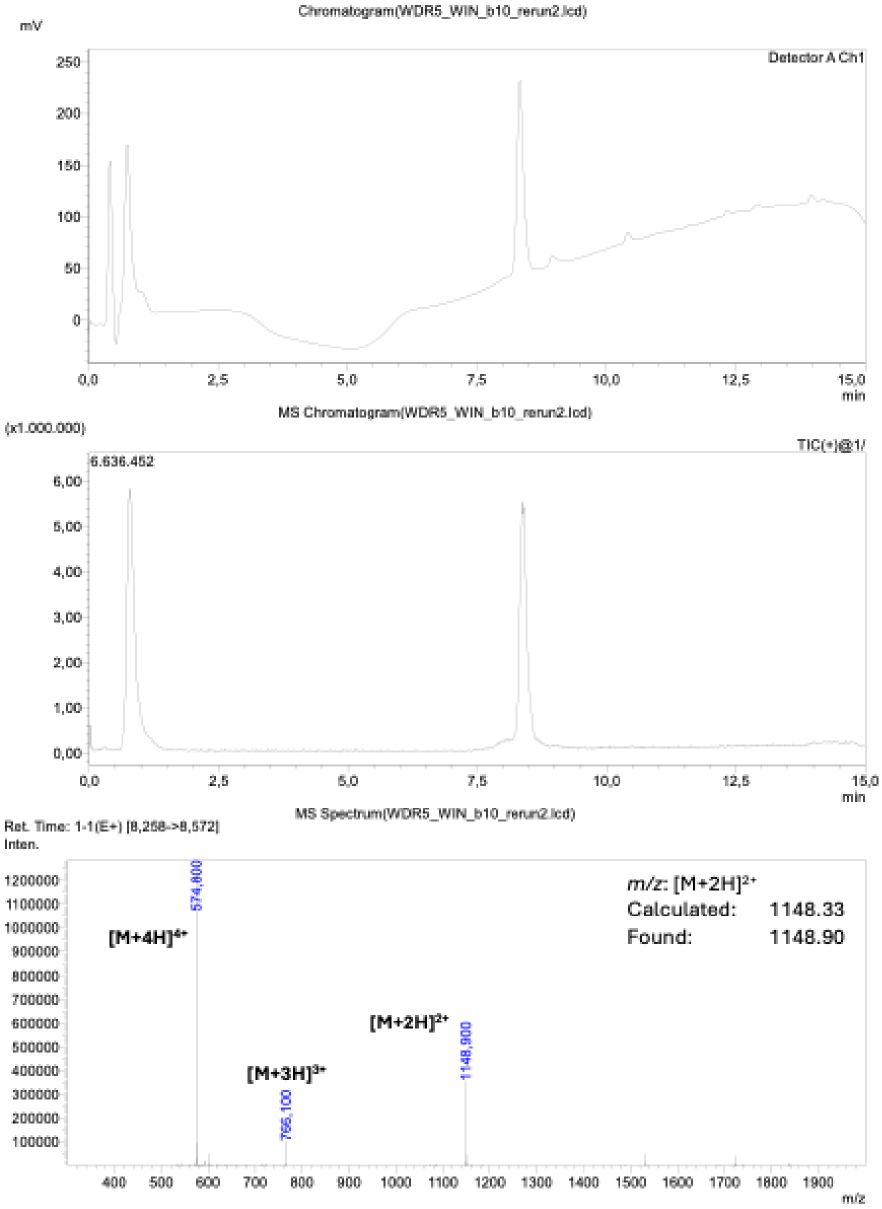

### Supporting Figure 32: LC-MS Analysis WDR5_WIN_B11

**WDR5_Win_B11:**
Ac-K(Biotin)-βA-SGKEQFEEMRNSMR-$CONH_2$

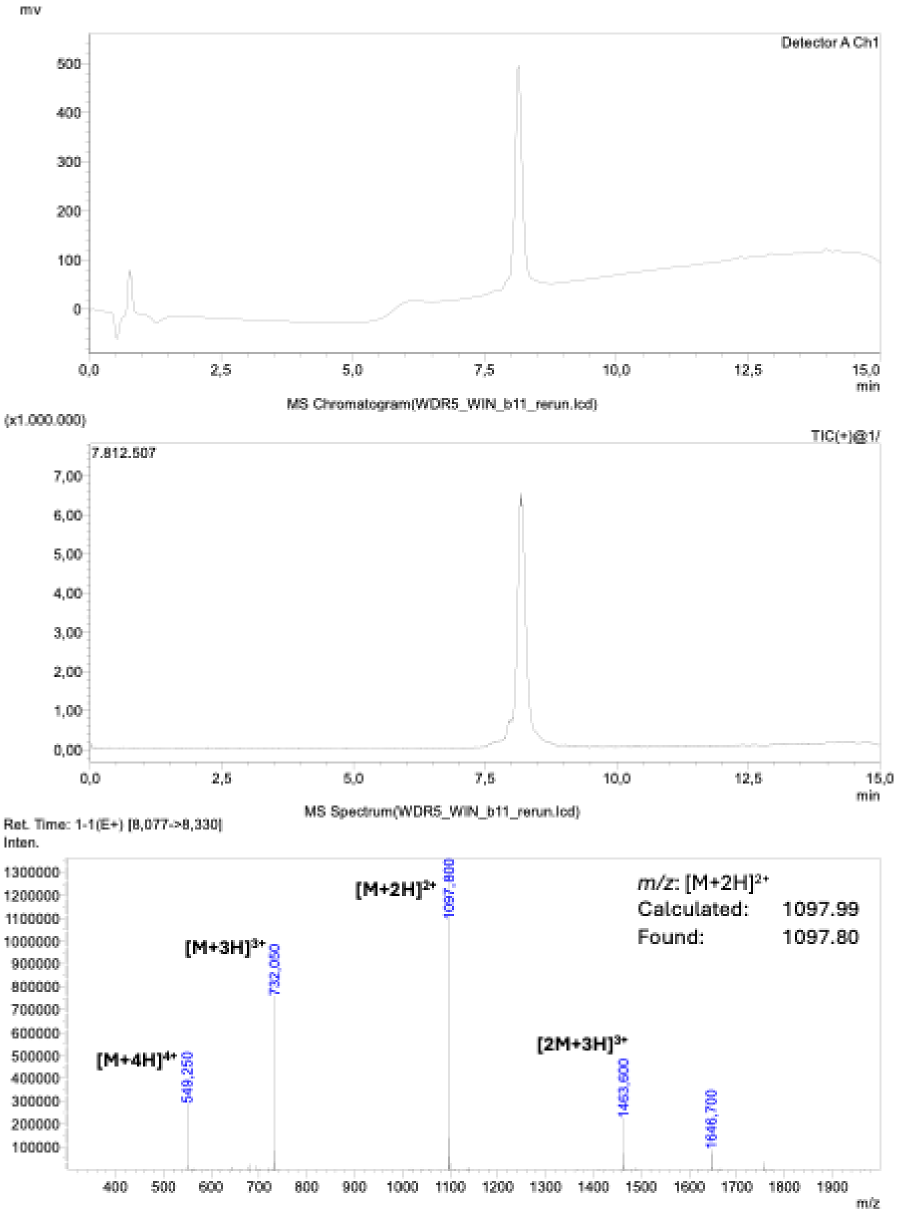

**Supporting Figure 33: LC-MS Analysis Myc_peptide_Motif.** Peptide used for WDR5-Myc competition assay.

**Myc-Peptide Motif:**
Ac-K(Biotin)-βA-EEIDVVSV-$CONH_2$

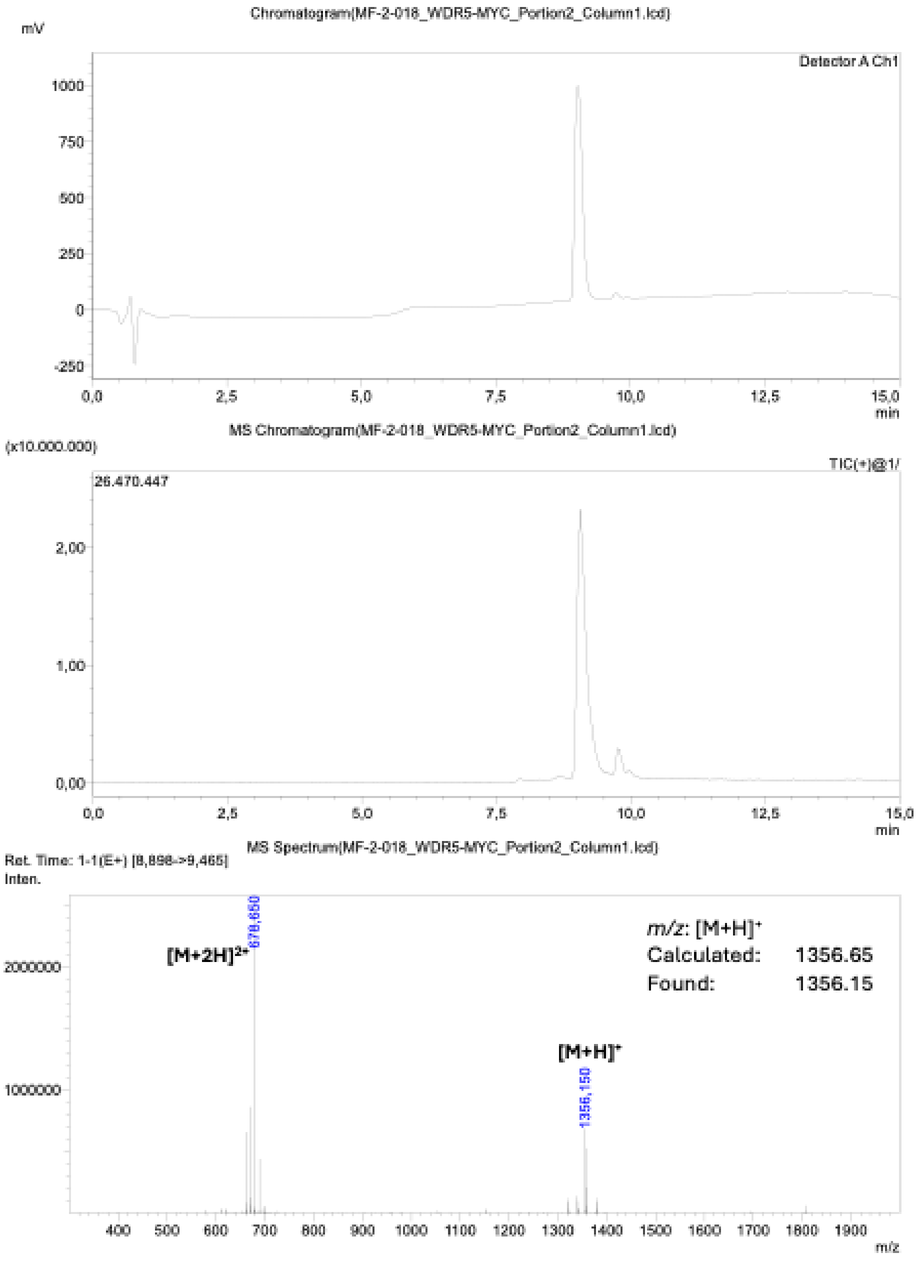

**Supporting Figure 34: LC-MS Analysis p53_peptide_Motif.** Peptide used for MDM2-p53 competition assay.

**MDM2-p53-Peptide Motif:**

Ac-K(Biotin)-βA-ETFSDLWKLLPE-$CONH_2$

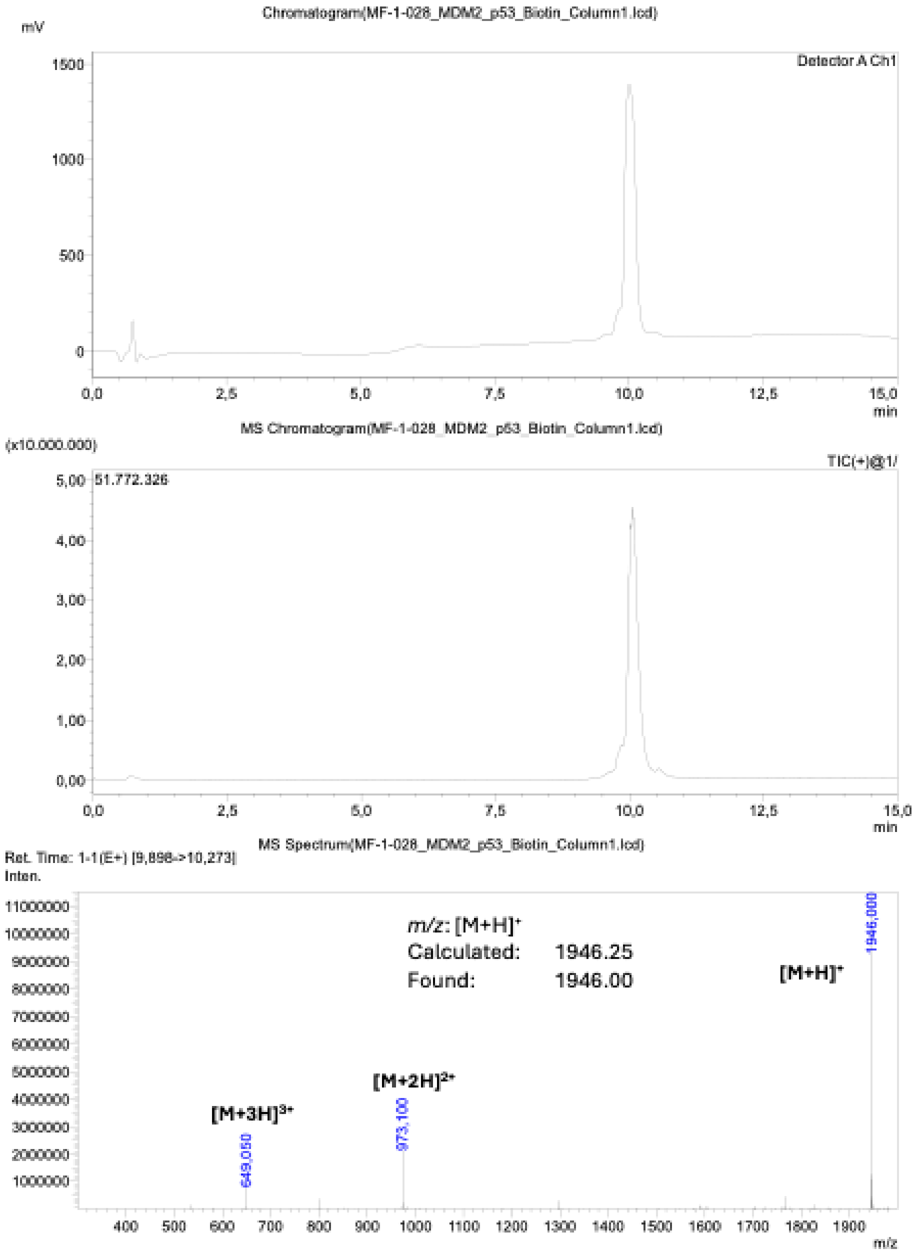

**Supporting Figure 35: LC-MS Analysis MDM2_B1_compeition peptide.** Peptide used for MDM2 competition assay.

**MDM2_B1 for competition:**
Ac-GQSWVNKFMEDMMS-$CONH_2$

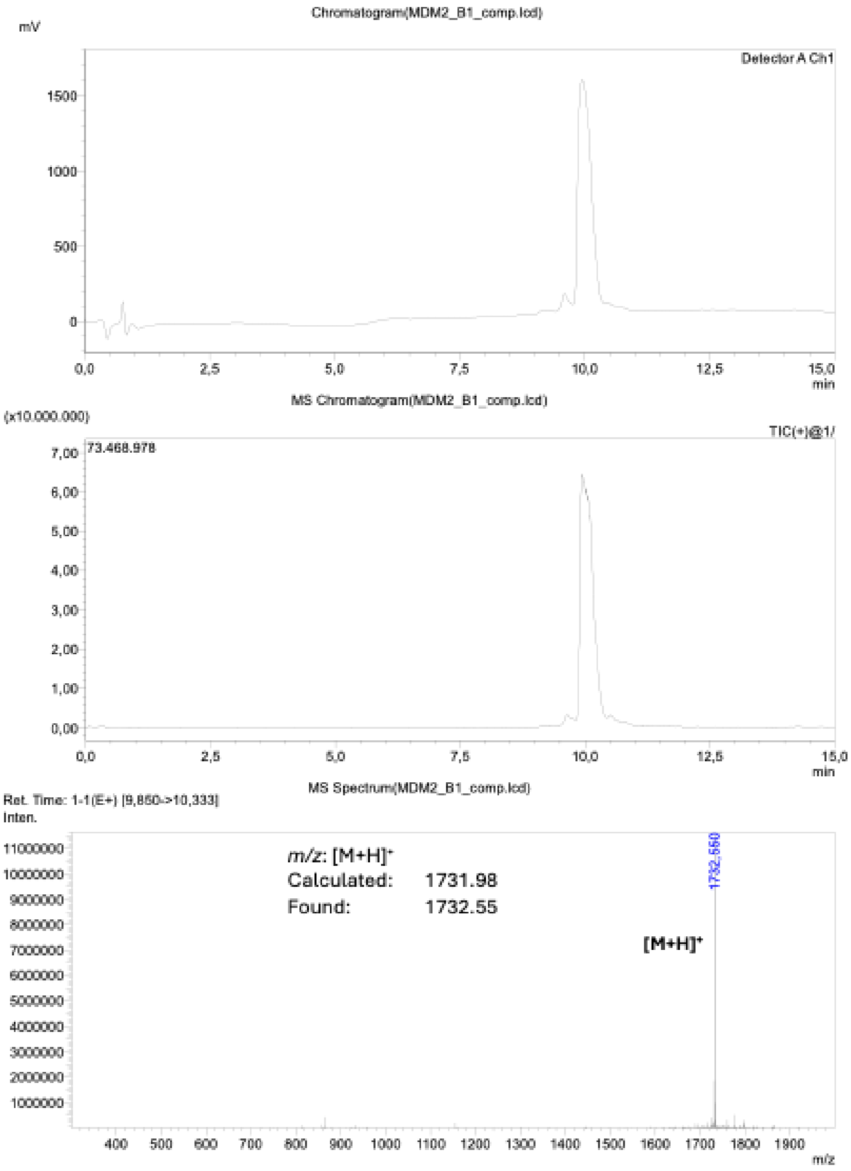

**Supporting Figure 36: LC-MS Analysis MDM2_B1_compeition peptide.** Peptide used for MDM2 competition assay.

**MDM2_B17 for competition:**
Ac-SPSEFQKHWQDLWDDYMK-$CONH_2$

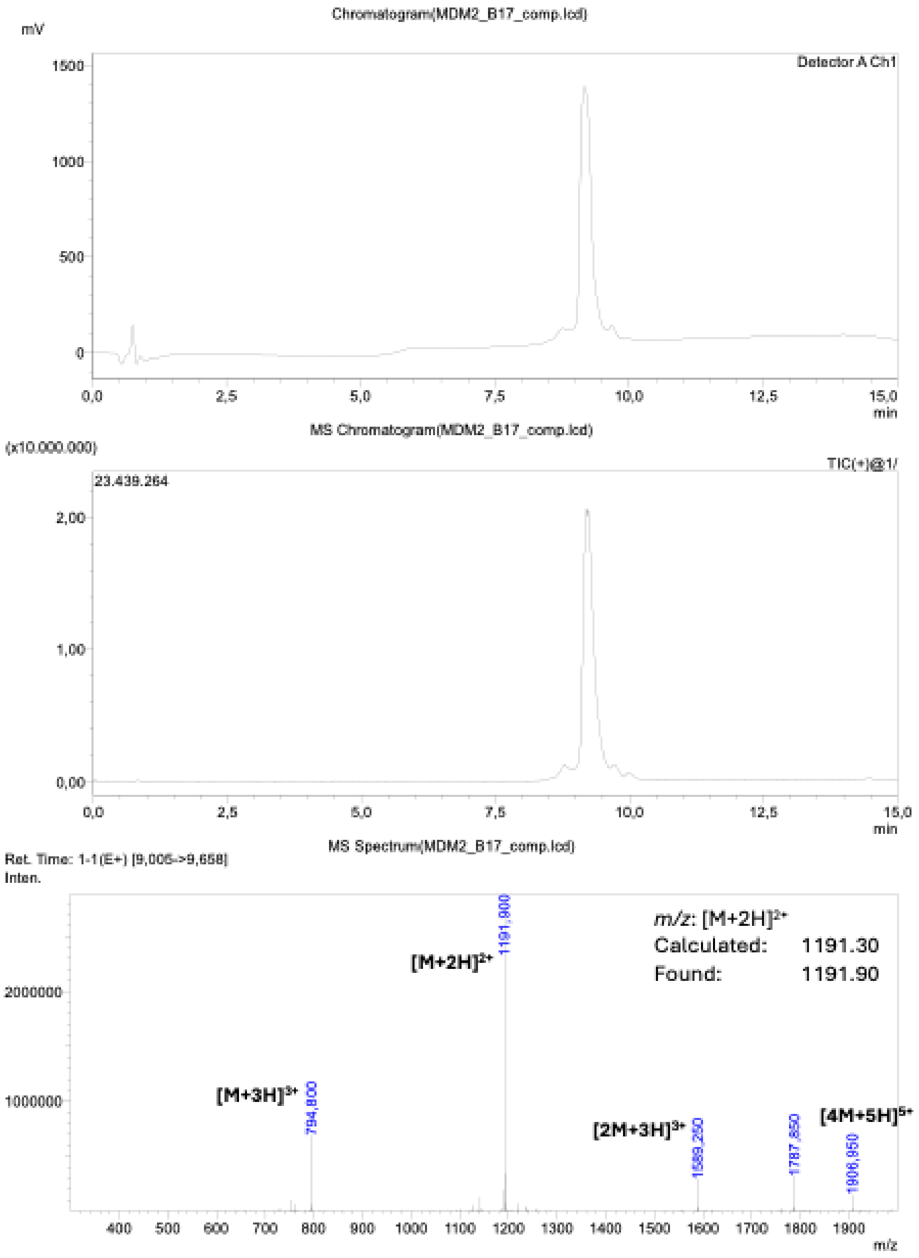

**Supporting Figure 37: LC-MS Analysis WDR5_Myc_B1_compeition peptide.** Peptide used for WDR5 competition assay.

**WDR5_Myc_B1 for competition:**

Ac-DDEDFEQFMKDLDEFLK-$CONH_2$

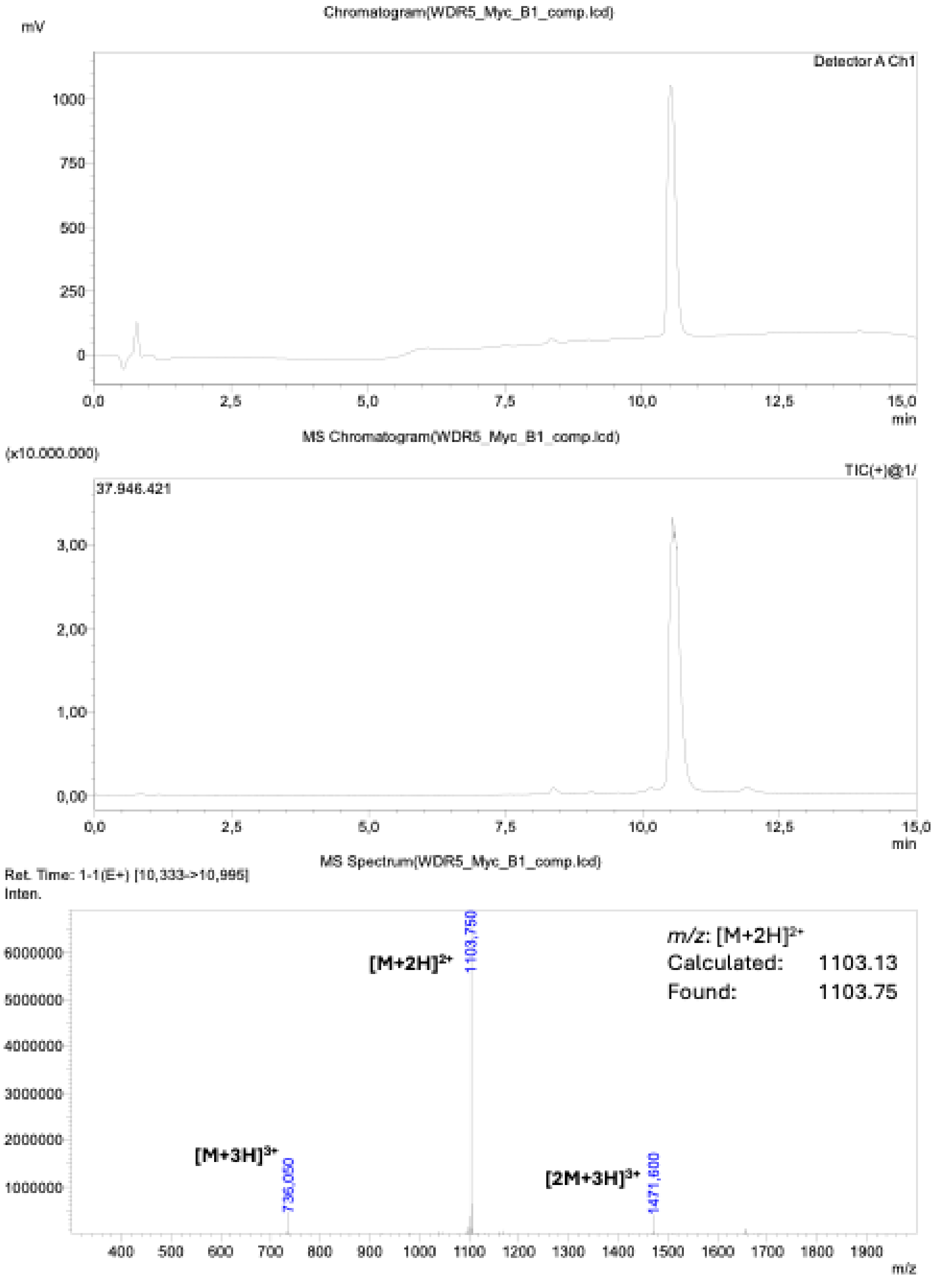

**Supporting Figure 38: LC-MS Analysis WDR5_Myc_B1_stapling_biotin.**

**WDR5_Myc_B1_biotin_stapled:**
Ac-K(Biotin)-βA-DDEDF**C**QFM**C**DLDEFLK-$CONH_2$

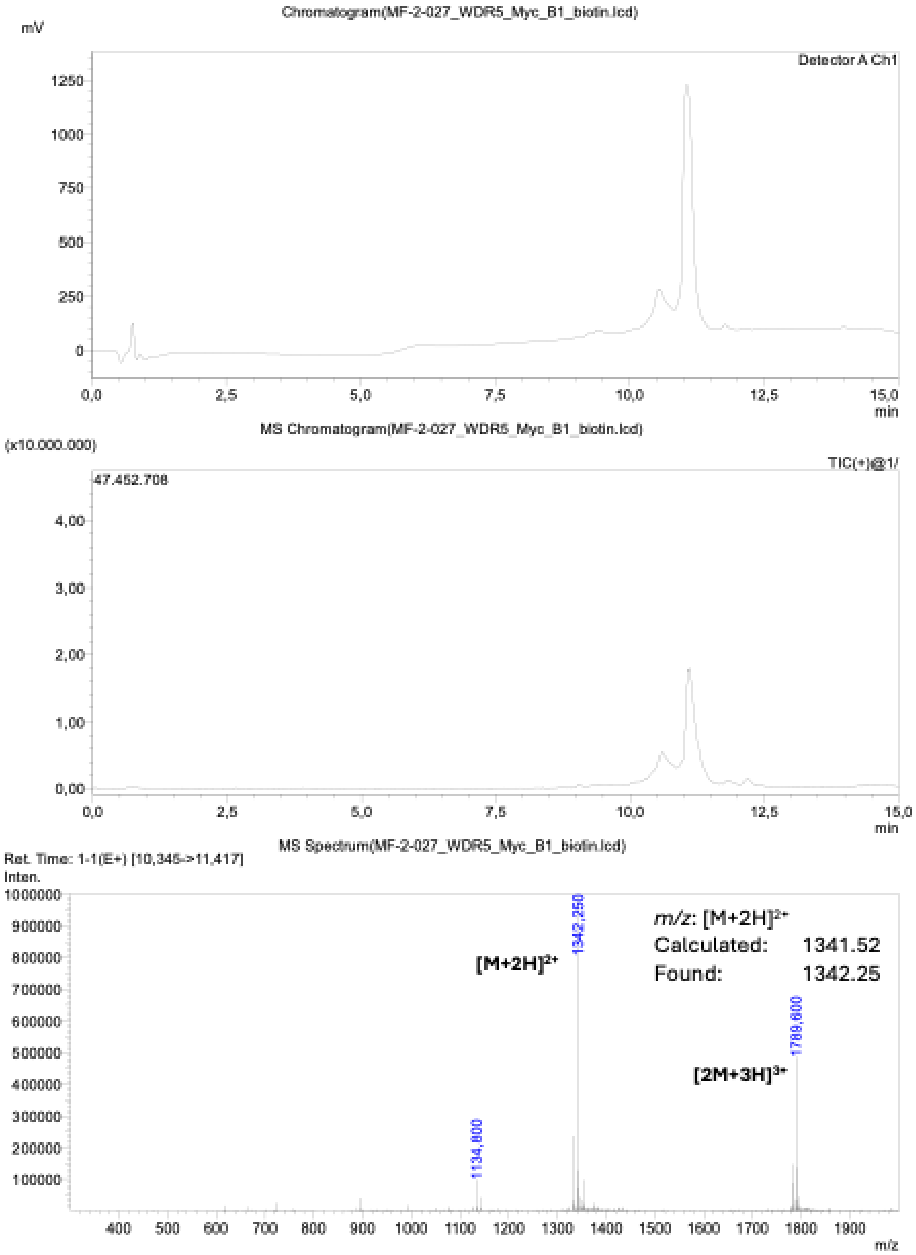

**Supporting Figure 39: LC-MS Analysis WDR5_Myc_B1_stapling competition assay.**

**WDR5_Myc_B1_NO_biotin_stapled_for_comp:**
Ac-βA-DDEDF**C**QFM**C**DLDEFLK-$CONH_2$

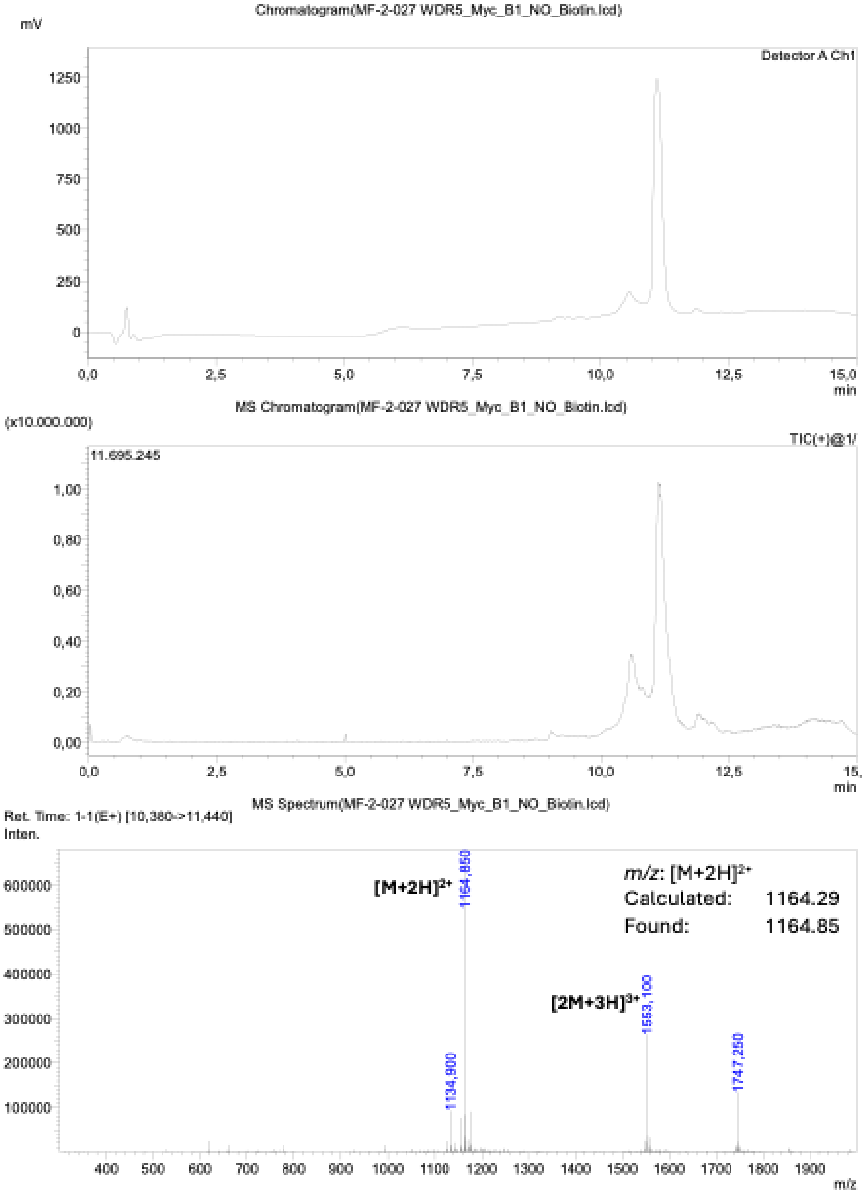
